# Pten is a disconnecting node in the molecular landscape of the proliferation/quiescence decision during mammary gland acinogenesis

**DOI:** 10.1101/2023.03.13.532403

**Authors:** Rebeka Tomasin, Ana Maria Rodrigues, Antonio Carlos Manucci, Alexandre Bruni-Cardoso

## Abstract

Cell context is key for cell phenotype. Using physiologically relevant models of laminin-rich ECM (lrECM) induction of mammary epithelial cell quiescence and differentiation, we have provided a landscape of the status of key molecular players involved in the proliferation/quiescence decision. Repression of some positive regulators of the cell cycle, such as cyclins and CDKs, occurred already at the mRNA level, whereas negative regulators of the cell cycle, such as Pten and p27, were upregulated only at the protein level. Interestingly, cell cycle arrest occurred despite the active status of Fak, Src and PI3k, because their downstream proliferative signalling pathways were repressed, suggesting the existence of a disconnecting node between upstream and downstream proliferative signalling in quiescent cells. Pten fulfils this role. Inhibition of Pten increased proliferation and restored Akt/mTORC1/2 and Mapk signalling in cells exposed to lrECM. In mice, Pten levels were positively correlated to the basement membrane thickness in the developing mammary epithelia, and Pten localized to the apicolateral membrane of luminal cells both in in ducts and near the nascent lumen in terminal end bud, characteristics consistent with a role for Pten in inducing and sustaining quiescence and tissue architecture. Accordingly, in 3D acininogenesis models, Pten was required for the onset and maintenance of quiescence, cell polarity and lumen assembly. The notion that lrECM-triggered differentiation involves a signalling circuitry with many layers of regulation provides an explanation for the resilience of quiescence within a growth-suppressive microenvironment, and that perturbations in master regulators, such as Pten, could disrupt the quiescent phenotype.

## Introduction

The mammary gland is unparalleled regarding to development and differentiation. A rudimentary epithelial tree at birth, the mammary gland further branches in puberty and will only fully grow and differentiate during pregnancy and lactation^1^. Upon weaning, the mammary gland goes through involution and massive remodelling, returning to a histological architecture similar to the one found before pregnancy^1^. This cycle repeats as many times as pregnancy, lactation and weaning occur in a female mammal lifetime^1,2^. All these events are coordinated by local and systemic stimuli^1^. Therefore, the developing mammary gland offers an invaluable tool for the study of spatiotemporal control of cell behaviour by the cell microenvironment.

A complex and widely diverse mesh of proteins and soluble factors, the ECM not only confers physical support for the cells, but also provides biochemical and mechanical cues perceived by the cells^1,3^, that respond accordingly altering cell phenotype, which in turn, remodel the ECM in a dynamic and reciprocal process^4^. Cells attach to specific ECM molecules via surface receptors, including integrins and the dystroglycan receptor^5,6^. Intracellularly, these receptors are physically linked to a range of cytoskeleton and signalling proteins, hence, mechanically and biochemically connecting cells to insoluble extracellular cues^3,6^.

Thus, the composition and mechanical properties of the ECM, along with the receptors and downstream effectors expressed by the cells will dictate cell behaviour, modulating processes such as proliferation, quiescence, differentiation, motility and polarity^3,4^.

Epithelial cells reside on top of a specialized ECM compartment, the basement membrane (BM), which formation and microarchitecture are nucleated and stabilized mainly by laminins^7^. The BM physically separates epithelial cells from the surrounding stroma, in which the ECM is mainly constituted by collagen I and fibronectin^7^. An intact BM is essential to sustain epithelial homeostasis^8,9^.

During pubertal mammary epithelial development, the BM is thicker in ducts than in TEBs^1^ and this has been extensively linked to the spatiotemporal regulation of mammary gland morphogenesis *in vivo* and in robust 3D cell culture models, where cells retain the ability to respond to laminin-triggered quiescence and differentiation, even in the presence of mitogens and growth factors^2,10–12^. A key event during epithelial morphogenesis is the rise of cell polarity, that along with cell-cell and cell-matrix interactions dictates tissue architecture^3^.

We demonstrate here that lrECM-triggered quiescence in mammary epithelial cells involved activation or suppression of several signalling pathways that control proliferation. Downregulation of some positive regulators of the cell cycle (cyclins and CDKs) occurred already at the mRNA level, whereas negative regulators of the cell cycle, such as Pten and p27, were upregulated only at the protein level. Interestingly, cell cycle arrest was achieved despite the active status of Fak, Src and PI3k, all positive regulators of proliferation. But even though Fak, Src and PI3k were active, their downstream effectors were not: Akt, mTORC1/2 and Mek/Erk were all found to be dramatically downregulated in quiescent cells, which could be an indicative of a “disconnecting node” between upstream and downstream proliferative signalling in laminin-induced quiescence. Our data pointed out that Pten fulfils this role.

Pten is a dual phosphatase that targets both proteins and lipids, but that also presents some non-enzymatic functions, particularly in the nucleus, where it promotes genome stability^13^. Pten is largely recognized by its tumour suppressor role counterbalancing Pi3k activity, where it hydrolyses PIP3 back to PIP2^14^. In addition to inhibiting proliferation, Pten has also been linked to other biological processes that are essential for epithelial morphogenesis and differentiation, such as cell polarity and lumen formation^15–19^.

We found that Pten was upregulated in laminin-rich ECM-treated mammary epithelial cells in 2D and its inhibition hindered quiescence by restoring the activity of PI3k/Akt, mTORC1/2 and Mek/Erk pathways. Correspondingly, the levels and tissue distribution of Pten correlated to the degree of laminin staining and the presence of lumen in the developing mammary gland. Pten levels were higher in ducts, which are laminin-rich, polarized, differentiated and mostly quiescent, than in TEBs, which are the laminin-poor, proliferative and invasive tips of the growing gland. In addition to higher levels, Pten was polarized to the apicolateral membrane in ducts, and curiously, also in cells near the nascent lumen in TEBs. Tests on mammary 3D acini, have shown that Pten inhibition increased cell proliferation in both forming and mature structures, disrupted polarity and led to loss of the lumen. Therefore, Pten acts a central node simultaneously coordinating differential, yet complementary events necessary for mammary gland development and homeostasis, including proliferation, polarization and lumen maintenance.

Altogether, these data support a model where laminin-induced quiescence was achieved by the modulation of a range of signalling pathways at different levels. Pten regulation seemed key for mammary gland development and homeostasis; with its downregulation being necessary for epithelial cell growth in the terminal end buds and its upregulation sustaining quiescence and tissue architecture in mature structures.

## Results

### Laminin-induced quiescence is characterized by alterations in cell size and morphology, G1 arrest and decreased CDK activity

When the basement membrane that envelopes the mammary epithelia is mimicked in culture systems using laminin-rich ECM (lrECM), primary epithelial cells and some established non-malignant epithelial lines, such as the murine EpH4 cells, become proliferative quiescent^20^, comprising a physiological and very suitable model for the study of microenvironmentally-regulated cellular quiescence.

As previously reported, lrECM triggered remarkable morphological alterations and G1 arrest in EpH4 cells^10,20,21^, reducing the overall cell number without affecting cell viability (Fig. 1 A-E). LrECM-induced G1 arrest was consistently observed either by analysing DNA content via flow cytometry (Fig. 1 D) or via fluorescence microscopy of EpH4-ES-FUCCI cells (Fig. 1 E). The FUCCI system is based on the cyclic degradation of fluorescent probes for Cdt1 (mCherry) and Geminin (mCitrine). Once Cdt1 accumulates during G1 and is degraded in early S phase, whereas Geminin builds up over S/G2/M, FUCCI cells express mCherry during G1 and mCitrine during S/G2/M, allowing for monitoring cell cycle in live cells^22^.

**Figure 1.**
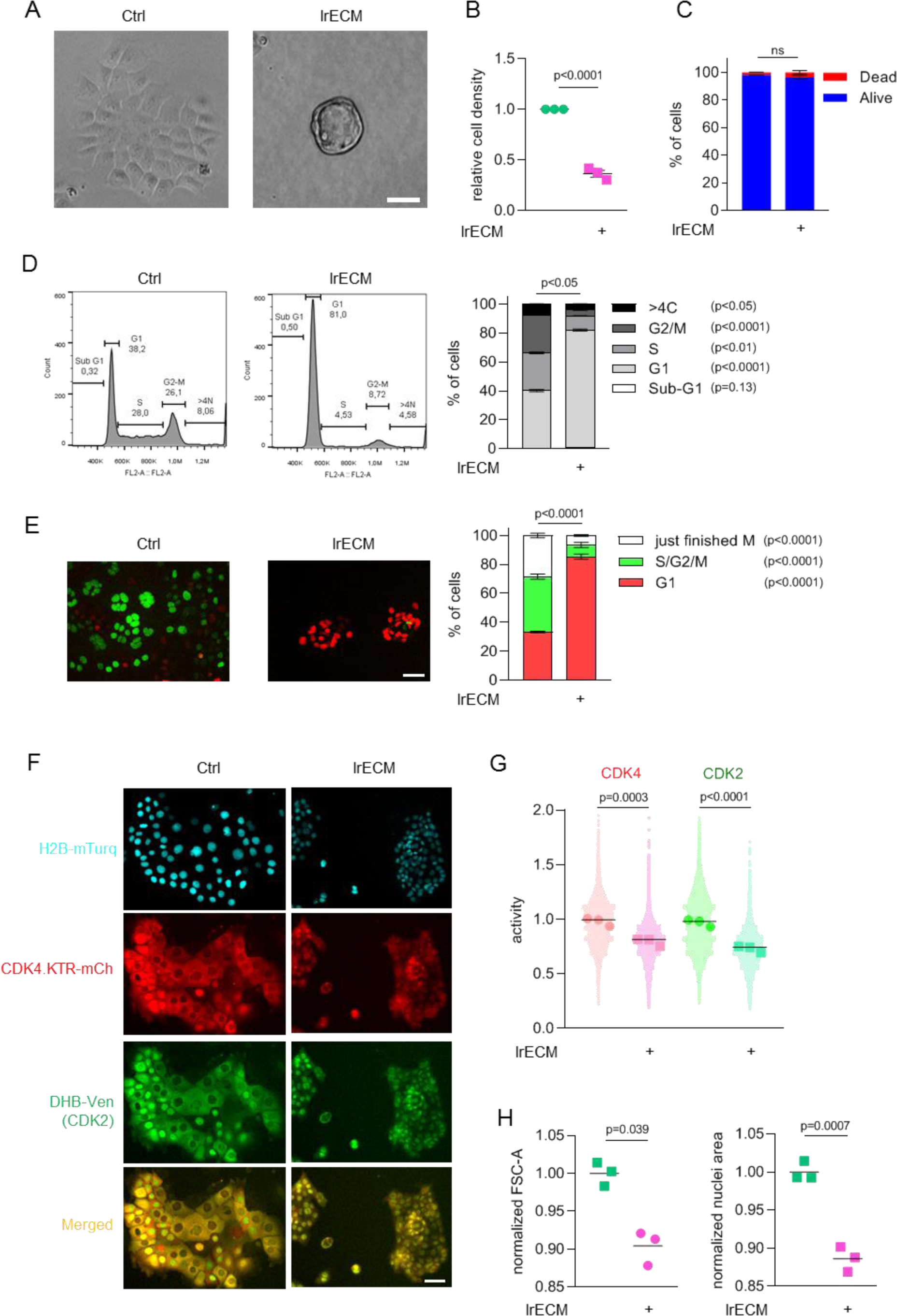
lrECM treatment of mammary epithelial cells induces G1 cell cycle arrest, reduces cell number and leads to morphological changes without inducing cell death. A-D EpH4 cells were treated or not with lrECM overlay for 48h. A. Bright field images of freely cycling (control) or quiescent (lrECM-treated) cells (scale bar 50 µm). B. Relative cell density (dots: normalized mean for each replicate, line: mean of all replicates, n=3, unpaired t-test, two tailed). C. Percentage of live and dead cells (mean±SEM, n=3, Two-way ANOVA followed by Sidak’s multiple comparisons test). D. Cell cycle analysis (PI staining & flow cytometry). Left: Representative histograms displaying count (Y axis) vs PI fluorescent intensity (X axis). Right: Distribution of cells across the cell cycle phases according to DNA content (sub-G1, G1, S, G2/M and >4C) in freely cycling (control) or quiescent (lrECM-treated) cells (mean±SEM, n=3, Two-way ANOVA followed by Sidak’s multiple comparisons test). E. Cell cycle analysis using live imaging of EpH4 cells expressing the ES-FUCCI sensor subjected to lrECM overlay assay for 48h (n=3 biological replicates, number of cells analysed per biological replicate: control=10010±429, lrECM=3567±409, mean±SEM, One way ANOVA followed by Sidak’s multiple comparisons test). Right: Fluorescent images of live EpH4-FUCCI cells freely cycling (control) or quiescent (lrECM). mCit+ cells (green) are in S/G2/M whereas mCherry+ cells (red) are in G1 (scale bar 50 µm). Left: Distribution of cells across the cell cycle according to the expression of the FUCCI sensors (G1, S/G2/M and just finished M/very early G1) in control or lrECM-treated cells (% mean±SEM). F-G. CKD4 and CDK2 activity in EpH4 cells upon lrECM overlay treatment for 48h (n=3 biological replicates, number of cells analysed per biological replicate: control=19658±3443, lrECM=6334±1008, mean±SEM). F. Fluorescent images of live EpH4-DHB-mVen-mCh-CDK4.KTR-H2B-mTurq cells freely cycling (control) or quiescent (lrECM-treated). Cyan: H2B-mTurquoise (nuclear marker); green: DHB-mVenus (CDK2 activity sensor); red: mCh-CDK4.KTR (CDK4 activity sensor); merged image: DHB-mVenus and mCh-CDK4.KTR (scale bar 50 µm). G. CDK4 and CDK2 activity (smaller dots: values for individual cells, bigger dots: mean of each biological replicate, line: mean). H. Cell size of cells in G1 treated or not with lrECM for 48h (dots: normalized mean for each replicate, line: mean of all replicates, n=3 biological replicates). Left: Cell size estimated by Flow cytometry (normalized FSC-A). Right: Cell size estimated by microscopy (normalized nuclear area).

LrECM-induced quiescence was also accompanied by decreased CDK4 and CDK2 activities (Fig. 1 F-G). CDK2 and CDK4 activities were evaluated using specific kinase translocation reporters (KTR) ^23,24^. In this system, cells express a nuclear marker (H2B-mTurquoise) and probes for CDK2 (DHB-mVenus) and CDK4 (mCherry-CDK4KTR) activities. Upon phosphorylation by CDK2 or CDK4/6, the fluorescent probes are translocated from the nucleus to the cytoplasm. Therefore, high CDK activity would result in the probe being translocated to the cytoplasm, and thus, CDK activity can be estimated by the ratio of cytoplasmic and nuclear mean fluorescent signal of a given probe^23,24^.

Quiescent cells were smaller than freely cycling cells, even considering only cells in G1 (Fig. 1 H). This was consistent when size was estimated via flow cytometry (Fig. 1 H, left) or via microscopy (Fig. 1 H, right).

Many molecular circuits could be involved in the acquisition and maintenance of the quiescent phenotype, what probably accounts for the morphological and functional diversity amongst quiescent cells in nature^25^. We sought to delineate a landscape of key molecules involved in cell cycle progression are regulated during laminin-induced quiescence in mammary epithelial cells.

### Laminin-induced quiescence and differentiation in mammary epithelial cells is accompanied by alterations in cell cycle regulators at the transcriptional and protein levels

In addition to becoming quiescent in the presence of lrECM, upon concomitant exposure to prolactin, EpH4 cells become functionally differentiated for lactogenesis and produce milk-related proteins^26^. Thus, the EpH4 model is largely used for the study of cellular and molecular mechanisms underlying mammary gland functional differentiation. In this sense, untreated (proliferating) cells could be paralleled to the proliferative terminal end buds found in the developing mammary gland, whereas lrECM-treated cells would be comparable to the quiescent cells found in the ducts and immature alveoli. Similarly, cells treated with both lrECM and prolactin would represent cells found in the alveoli of fully mature, milk-producing mammary gland.

To obtain a molecular landscape of quiescence in the context of lrECM, we compared proliferating, quiescent and lactogenic EpH4 cells regarding some key cell cycle regulators, initially at the transcriptional level (Fig. 2).

**Figure 2.**
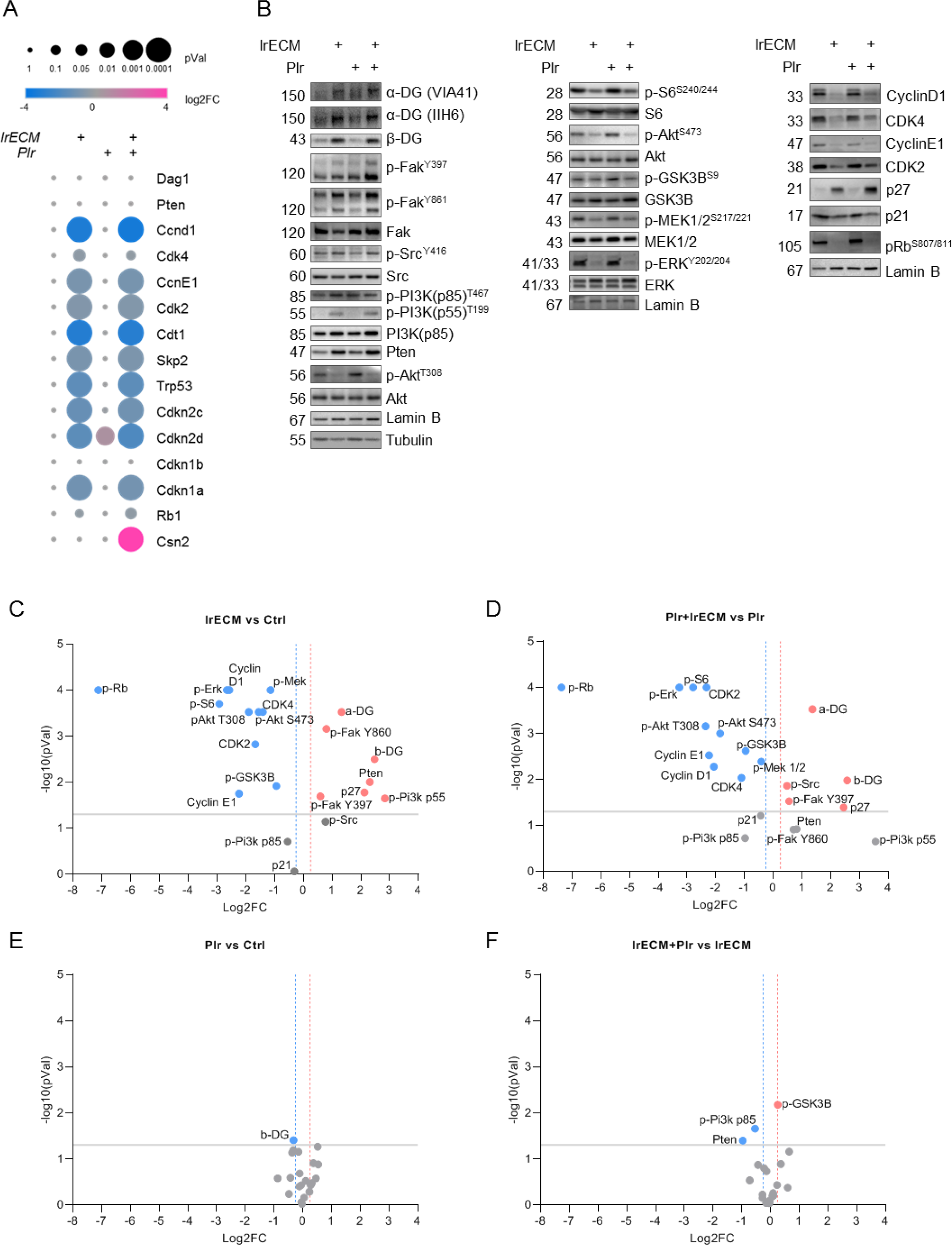
Quiescent epithelial cells sustain the activity of Fak, Src and Pi3k but display decreased signalling in downstream positive regulators of cellular proliferation. EpH4 cells were treated or not with lrECM overlay and/or prolactin (Plr) for 48h and evaluated for mRNA and protein expression levels of some key cell cycle regulators (n=3). A. Relative mRNA expression levels assessed by RT-qPCR (One way ANOVA, followed by Dunnett’s multiple comparison test comparing all samples to the Control). B. Immunoblotting of key proliferation and cell cycle regulators for protein levels assessment. Left panel: Upstream regulators of cell proliferation. Mid panel: Intermediate mediators of cell proliferation. Right panel: Downstream positive and negative regulators of cell cycle. C-F Volcano plots displaying direct comparisons between two samples (logFoldChange vs –log10pValue). Comparison between two groups were performed by unpaired t-test (two tailed), p<0.05 (-logpval = 1.30) was considered significant. Dashed lines mark a 25% decrease (blue) or increase (red). C. lrECM-treated (quiescent) vs Ctrl (proliferative) cells. D. Cells treated with lrECM+Plr (quiescent and lactogenic) vs cells treated only with Plr (proliferative). E. Plr (proliferative) vs Ctrl (proliferative) cells. F. Cells treated with lrECM+Plr (quiescent and lactogenic) vs. lrECM-treated (quiescent) cells.

The mRNA levels of several cell cycle regulators dramatically changed upon lrECM treatment, regardless of cotreatment with prolactin (Fig. 2A). Confirming the functional differentiation state towards a lactogenic program^26^, only the exposure to both lrECM and prolactin triggered the expression of *Csn2*, the gene encoding β-casein (Fig. 2A). The mRNA expression of most positive regulators of the cell cycle (such as Cyclins and Cdks) plummeted upon lrECM treatment (Fig. 2A), but, surprisingly, so was the expression levels of some negative regulators of cell cycle (such as *Trp53*, *Cdkn1a*, *Cdkn2c and Cdkn2d*). On the other hand, the mRNA levels of *Pten*, *Dag1* and *Cdkn1b* were unchanged across all treatments (Fig. 2A). These data confirm some previous findings indicating that upon laminin-induced quiescence acquisition there is a general repression of transcription^10^. But how are these cells arrested if, apparently, negative regulators of the cell cycle were either unchanged or also down? The lack of alteration or even the decrease observed in mRNA levels of cell-cycle negative regulators might indicate that these molecules are most likely to be controlled at the protein level during lrECM-induced quiescence.

Indeed, many of the alterations in cell cycle regulators happen at the protein level^27^ and therefore, we have assessed the protein levels of a range of known positive and negative regulators of the cell cycle by western blot (Fig. 2B-F).

lrECM-induced quiescent cells displayed dramatic changes in protein expression and/or phosphorylation levels of key cell cycle regulators (Fig. 2B-F). Again, lrECM-treated cells exhibited similar protein expression profiles concerning cell cycle regulators, regardless of prolactin supplementation (lrECM *vs* lrECM+Plr - Fig. 2B, F), and, likewise, cells treated only with prolactin exhibited a protein expression profile similar to untreated cells (Fig. 2B, E).

Quiescent cells displayed decreased protein levels of positive regulators of proliferation, such cyclins and CDKs (Fig. 2B, right panel, C and D), reflecting their transcriptional repression (Fig. 2A), which resulted in the almost absence of hyperphosphorylated Retinoblastoma protein (Rb) in lrECM-treated quiescent cells (Fig. 2B, right panel, C and D). Complementarily, quiescent cells also had increased levels of some negative regulators of the cell cycle, such as Pten^28^, p27^29^ and the dystroglycan receptor^30^, which had shown no alterations at the transcript level (Fig. 2A). Once the balance between cyclins and CKIs is believed to be one of the components that dictated cell cycle progression^31,32^, the decreased activity of CDK4 and CDK2 observed in lrECM-treated cells (Fig. 1 F, G) could be, at least in part, attributed to the lower levels of cyclins and CDKs and increased levels of p27 found in quiescent cells (Fig. 2A, B-D).

LrECM treatment attenuated many proliferative signalling pathways, such as the Mapk pathway^11^ (evidenced by reduced levels of phospho-Mek and phospho-Erk) and the PI3k/Akt/mTORC1/2 pathway^11^ (evidenced by reduced levels of phospho-Akt and phospho-S6) (Fig. 2 B-D).

Surprisingly though, lrECM-treated quiescent cells displayed increased or unchanged levels of active (phosphorylated) Fak, Src and PI3k (Fig. 2B-D). However, downstream signalling triggered by these pathways was widely downregulated in the presence of lrECM (Fig 2B-D). These included the PI3k/Akt/mTORC1/2 axis and the MAPK pathway (Fig. 2B-D).

Fak/Src/PI3k are components of adhesion complexes and are activated upon integrin binding to ECM molecules^33^. Fak, Src and PI3k also take part in the control of mitogenic signalling triggered by the binding of growth factors to RTKs^34^. There is significant cross-activation between Fak, Src and PI3k, and although each of them do have specific functions on cell behaviour, their activation usually triggers Akt/mTORC1/2 and MAPK signalling^35^, which places them as positive regulators of cell proliferation.

Decreased phosphorylation levels of Akt and Mek/Erk have been previously reported in EpH4 acini^36,37^ and also in non-malignant human mammary epithelial S1^11,38,39^ and MCF 10A^40,41^ cells undergoing laminin-induced quiescence.

The Ras/Raf/Mek/Erk axis is the most crucial MAPK signalling pathway for the positive regulation of cell proliferation in epithelial cells^42^. It is manly triggered by the binding of mitogens and growth factors to RTKs, but it is also modulated by the ECM^34,35,42^. Complementarily, mTOR controls cell growth by promoting anabolism and suppressing catabolism and its function might be different depending on upstream signalling^43^. Consistent with the smaller cell size observed in quiescent cells, we observed decreased phosphorylation levels of S6 and AKT^S473^, respectively reflecting the activities of mTORC1 and mTORC2 (Fig. 1H).

Another feature of quiescent cells that have captured our attention was the presence of phospho-PI3k p55 only in lrECM-treated cells (Fig. 2B). The PI3k complex is comprised of catalytic (p110) and regulatory (p85 or its alternative spliced versions p50 and p55) subunits. Specific functions for different PI3k isoforms in mammary gland development have been described, including a role of p110γ in lumen formation^44^ and for p55α/p50α for cell death induction during mammary gland involution^45^. Thus, phospho-PI3k p55 might have functions beyond the regulation of proliferation in lrECM-induced quiescence.

### Contribution of specific signalling pathways for cell cycle progression and the role of Pten as a disrupting node for Fak, Src and PI3k proliferative signalling during lrECM-induced quiescence

The data on protein expression prompted us to assess the contribution of different signalling pathways for the cell cycle arrest upon lrECM treatment by using commercially available inhibitors (Table 3,Fig. 3).

**Figure 3.**
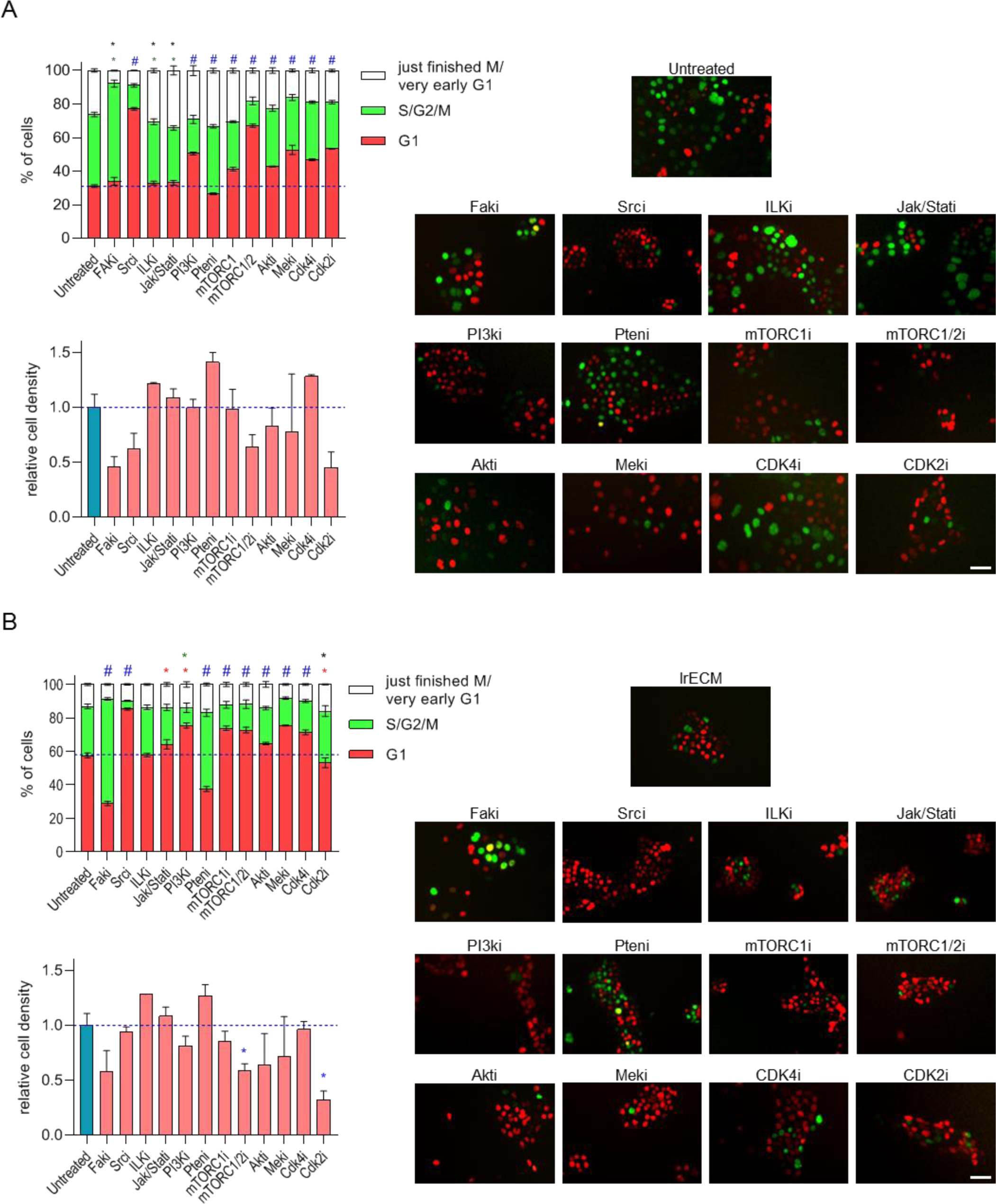
Inhibition of upstream and downstream regulators of the cell cycle affects the balance between proliferation and quiescence in mammary epithelial cells. FUCCI-based cell cycle distribution in EpH4-FUCCI cells treated or not with lrECM and specific inhibitors for 24h and representative images displaying mCherry (G1) and mCitrine (S/G2/M) positive cells for each condition (scale bar 50 µm). A. Cells treated only with the inhibitors. B. Cells treated with lrECM and inhibitors simultaneously. Experiments were performed in two biological replicates with two replicate wells per per biological replicate. Relative cell density was analysed by One way ANOVA followed by multiple comparisons by Two-stage linear step-up procedure of Benjamini, Krieger and Yekutieli (Ctrl vs different treatments), where q< 0.01 was considered significant. FUCCI-based cell cycle analysis was performed using Two-way ANOVA followed by multiple comparisons by Two-stage linear step-up procedure of Benjamini, Krieger and Yekutieli (Ctrl vs different treatments, for each cell cycle phase), where q< 0.01 was considered significant. # (blue) differences between G1, S/G2/M and just finished M/early G1 were all statistically significant. * (red) only the difference between G1 was statistically significant. * (green) only the difference between S/G2/M was statistically significant. * (black) only the difference between just finished M/early G1 was statistically significant. q<0.05 was considered significant. Fak: FAK inhibitor 14 (1 µM); Src: PP2 (2 µM); ILK: CPD22 (0.25 µM); Jak/Stat: Tofacitinib (20 µM); PI3K: LY294002 (20 µM); Pten: bpV(pic) (2.5 µM); mTORC1: Rapamycin (0.2 µM); mTORC1/2: Torin 2 (0.4 µM); Akt: MK 2206 (2 µM); Mek: PD98059 (20 µM); Cdk4: Palbociclib (1 µM), Cdk2: CDK2 inhibitor III (60 µM).

As expected, treatment with lrECM increased the percentage of cells in G1 (quiescent), and indeed, treatment with pharmacological inhibitors of intermediate and downstream positive regulators of the cell cycle (such as mTORC1/2, Akt, Mek, Cdk4 and Cdk2) mostly decreased cell cycle progression, and, in general, increasing the percentage of Eph4-ES-FUCCI cells arrested in G1 in both lrECM and Ctrl contexts (Fig 3A and B).

Despite the fact that both CDK2-CyclinE and CDK4-CyclinD1 were less active and had lower expression levels in quiescent cells (Fig. 1F, G,Fig. 2 A-D), the effects of CDK2 inhibition on cell cycle and cell number were more prominent than those observed for CDK4 inhibition (Fig. 3 A). Cells can often bypass loss of cyclins or CDKs and progress through cell cycle by activating alternative cyclin-CDK complexes^46–48^. Particularly, mature cyclin D-CDK4 complexes are refractory to Palbociclib, the CDK4 inhibitor used in this work. Palbociclib is only functional if binding to CDK4 by itself, before CDK4 complexing with cyclin D, and therefore inhibits cell cycle progression by preventing the assembly of new cyclin D-CDK4 complexes and their activity, and also by indirectly inhibiting CDK2^49^. Furthermore, the smooth-edged and compact morphology of cell clusters treated with CDK2 inhibitor (Fig. 3A, CDK2i) closely resembled quiescent lrECM-treated cells (Fig. 3B, lrECM), which was not the case for cells treated with the CDK4 inhibitor (Fig 3 A, CDK4i). These data could be an indicative that inhibition of CDK2 and CDK4 might have distinct roles in lrECM-induced quiescence.

Inhibition of Src, PI3k, mTORC and Akt all gave rise to small, G1-arrested cells, whereas Fak inhibition arrested cells in G2 (Fig 3A). Inhibition of Jak/Stat or Ilk had no or minor effects on cell cycle (Fig 3A). This is consistent with previous reports which have proposed a role for Jak/Stat signalling not for proliferation, but for milk production instead^26^. Similarly, Fak, Akt and Ilk signalling have all been linked to cell survival and/or polarity ^50,51^, whereas Src and PI3k/mTORC/Akt signalling are involved in the control of cell volume and size^43,52^.

Another key observation from this assay was that, despite the high levels of phosphorylated Fak, Src or PI3k proteoforms found in quiescent cells (Fig. 2), inhibition of these molecules did change cell proliferation (arresting cells in G1 for Src and PI3k inhibition and in G2 for Fak inhibition) (Fig. 3). Therefore, the activity of these signalling pathways, as a default, contribute for proliferation in mammary epithelial cells.

How could Fak, Src and Pi3k be active, but their downstream proliferative pathways be down in quiescent, lrECM-treated cells? There must be a “disconnecting point” between upstream and downstream proliferative signalling triggered lrECM treatment.

Pten, which was found upregulated in quiescent, lrECM-treated cells (Fig. 2B-D) could fulfil this role. Its lipid phosphatase function counterbalances PI3k activity by converting PIP3 to PIP2^28^, whereas its protein phosphatase activity can target Fak^53^.

Strikingly, Pten inhibition led to increased proliferation, increasing cell number and decreasing percentage of cells in G1/increasing percentage of cells in S/G2/M (Fig. 3). Alterations induced by Pten inhibition were clearer in lrECM-treated cells, where the distribution of cells across G1/S/G2/M resembled what was observed for freely cycling (control) cells (Fig. 3A and B).

Furthermore, inhibition of Pten partially restored PI3k (phospho-Akt^T308^), MAPK (phospho-Erk1/2), mTORC1 (phospho-S6) and mTORC2 (phospho-Akt^S473^) signalling in lrECM-treated cells (Fig. 4). Treatment with Pten inhibitor in the absence of lrECM had overall milder effects in further increasing phosphorylated levels of Fak, Akt, Erk and S6 (Fig 4), which goes along with the notion that subconfluent freely cycling (control) cells in culture are in exponential growth, near the maximum of their proliferation rates^54^.

**Figure 4.**
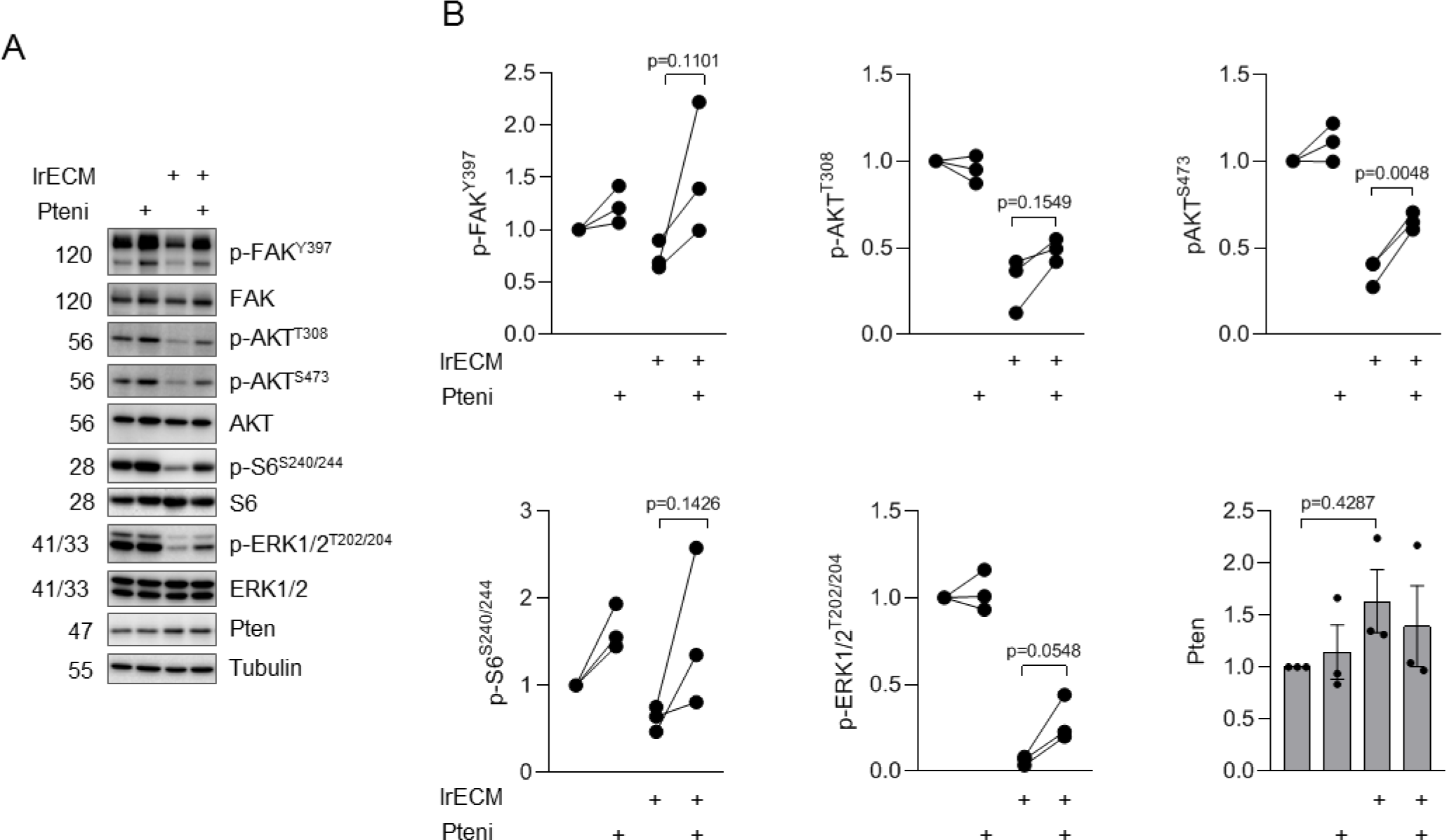
Inhibition of Pten reverts lrECM-induced quiescence by partially restoring PI3K, ERK, mTORC1 and mTORC2 signalling. A. Immunoblots of EpH4 cells treated or not with lrECM overlay and/or Pten inhibitor (bpV(pic) 2.5 µM) for 24h (n=3). B Quantifications of blots displayed in A (dots: normalized expression of each biological replicate; connecting lines: link samples from a given replicate). phospho-Fak^Y397^ (Pten target); phospho-Akt^T308^ (PKD1 substrate); phospho-Akt^S473^ (mTORC2 substrate); phospho-S6^S240/244^ (mTORC1 substrate); phospho-Erk^Y202/204^ (Mapk pathway). Statistical differences were assessed by One way ANOVA followed by Tukey post test for multiple comparisons.

These data, in combination with the protein expression analysis (Fig. 2B-D), indicated that Pten could act as a disrupting node for Fak/Src/PI3k proliferative signalling in mammary epithelial cells, especially upon quiescence induction by a laminin-rich microenvironment.

### Pten correlates with laminin deposition in the developing murine mammary gland and displays distinct distribution and polarization patterns in ducts and in TEBs

Our data obtained for mammary epithelial cells pinpointed Pten as upregulated upon laminin treatment and as an important regulator of laminin-induced quiescence. In order to evaluate whether this evidence holds true *in vivo*, we evaluated the correlation between laminin deposition and Pten staining in developing murine mammary glands (Fig. 5). Ducts, which are laminin-rich, mostly quiescent epithelial structures in the tubuloalveolar mammary network^55^ had higher levels of Pten (Fig. 5B-C). TEBs, which are the laminin-poor, invasive and proliferative tip of the growing epithelia in the mammary gland^55^ had overall lower levels and an unclear pattern of Pten staining (Fig. 5B-C). Furthermore, Pten intensity was found to be positively correlated to the degree of laminin deposition (Fig. 5B, D).

**Figure 5.**
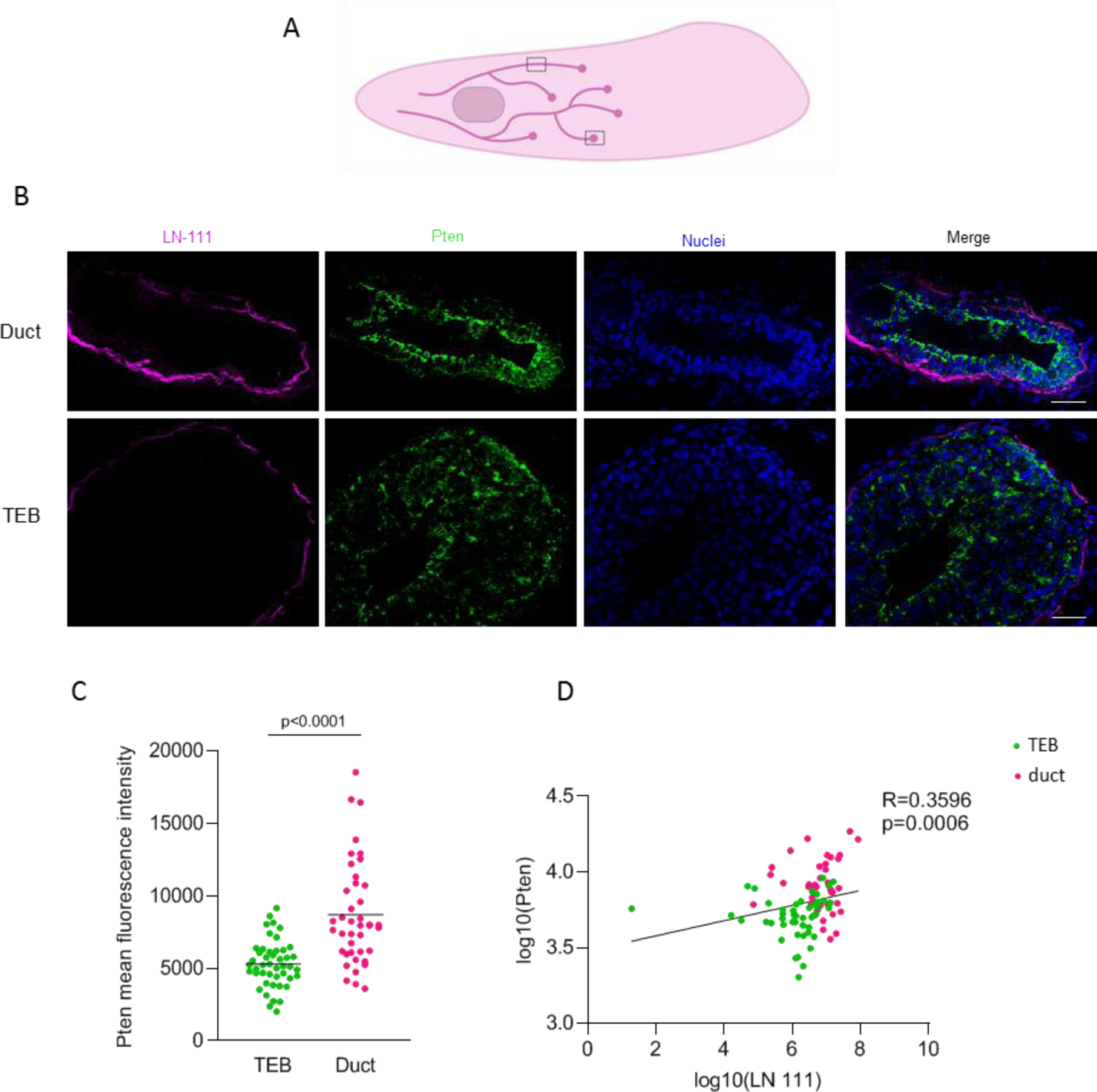
Pten levels are positively correlated with LN111 deposition in the developing mammary gland. Immunofluorescence analysis of Pten and Laminin levels in mammary glands of 5-weeks old female mice (n=3 mammary fat pads from 3 different mice). A. Schematic view of a developing mammary gland from a 5-weeks old female mouse. An example of areas corresponding to a duct (top) and a terminal end bud (TEB, bottom) are highlighted by dashed rectangles. B. representative images of a duct (top) and a TEB (bottom) stained for Pten (green) and laminin-111 (magenta) (scale bar 25 µm). C. Mean fluorescent intensity of Pten in cells found in TEBs (green) or ducts (pink). Statistical differences were evaluated by unpaired t-test (two tailed). Each dot corresponds to the mean of the mean fluorescence intensity of Pten per cell in a given field and the line marks the mean fluorescence intensity of Pten for all fields analysed in each group (ducts or TEBs). D. Correlation analysis (Spearman) of Pten levels and Laminin-111 deposition in TEBs (green dots) and ducts (pink dots). Data were plotted as log10. For further details on data analysis, please refer to the “Material and methods” section.

One feature that became clear while analysing the images of the developing mammary glands was that Pten presented distinct staining patterns in ducts and TEBs and amongst specific regions or cell types. Whereas in the ducts Pten showed strong apico-lateral staining in the membrane of luminal cells (Fig. 5B, top); this pattern was only seen in cells surrounding the nascent lumen in TEBs (Fig. 5B, bottom).

Once we observed a predominance of apicolateral Pten staining, particularly in polarized luminal epithelial cells in the duct, we next stained developing mammary glands for Pten and f-actin and imaged ducts and TEBs optically sliced at different focal planes (Z axis) using confocal and super resolution microscopy (Fig. 6). In ducts, Pten appears in puncta at the basal region (portion of the cell laying on the BM) and at the membrane along with the actin belt in the apical region (facing the lumen) (Fig. 6A). In TEBs (Fig. 6B), which mostly lack polarization, there was not a clear pattern of Pten staining. The exceptions were cap cells, found at the front edge of TEBs, where Pten was mainly found in the nuclei (Fig. 6B, mid panel) and of cells surrounding the nascent lumen, which presented a staining pattern of Pten and f-actin similar to what was observed in the apical region of ducts (Fig. 6B, top panel).

**Figure 6.**
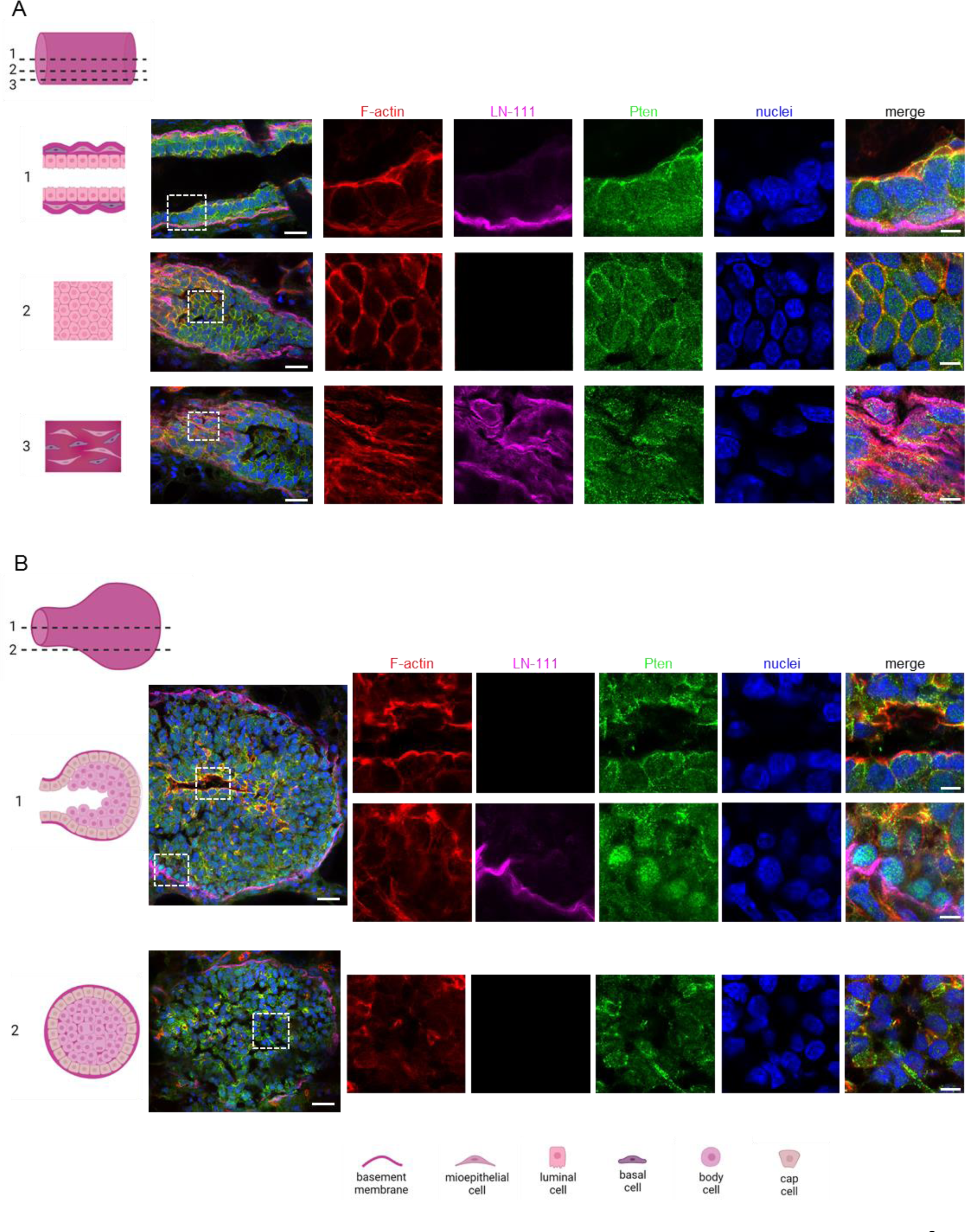
Pten colocalizes with f-actin in the apicolateral surface of luminal cells in the ducts and near the nascent lumen of TEBs. Confocal and super resolution microscopy images of epithelial cells in the developing mammary gland stained for f-actin (red), LN-111 (magenta), Pten (green) and counterstained with DAPI (blue) (n=3 mammary fat pads from 3 different mice). A. Duct. Left: Schematic representation of a duct, displaying to which position (Z) in the structure the following images correspond to (dashed lines and corresponding images 1-3) 1=central, 2=near the lumen (apical domain of luminal cells), 3=near the basement membrane (basal). The merged image displayed on the left was taken with a confocal microscope (scale bar 20 µm) and the dashed rectangles highlight the areas selected for Airyscan super resolution imaging (images with separated channels displayed on the right, scale bar 5 µm). Note apical Pten apical staining in images from positions 1 and 2. B. TEB. Left: Schematic representation of a TEB, displaying to which position (Z) in the structure the following images correspond to (dashed lines and corresponding images 1-2). 1=central, 2=closer to the basement membrane. The merged image displayed on the left was taken in a confocal microscope (scale bar 20 µm) and the dashed rectangles highlight the areas selected for Airyscan super resolution imaging (images with separated channels displayed on the right, scale bar 5µm). In the TEB (position 1), note apical Pten staining in cells near the nascent lumen (top images) and nuclear Pten staining in cap cells (bottom images). Images for different heights were not necessarily taken in the same field/structure.

The presence of Pten in the membrane is directly related to its phosphatase activity^56^ and might therefore indicate that Pten activity is not only higher in ducts and near the nascent lumen, but that this activity is predominantly polarized to the apical surface. The fact that cap cells, which are believed to be multipotent mammary stem cells ^57,58^, presented high Pten levels in the nuclei might be related to the contributions of Pten for genomic stability^13^, which would be particularly important for progenitor cells, once mutations in these would be passed on to all their descendant cells.

### Pten inhibition leads to cell proliferation and loss of cell polarity in both developing and mature mammary epithelial 3D acini

The distinct levels and cell and tissue localization of Pten in ducts and in TEBs prompted us to use cell culture 3D acini models to evaluate the contribution of Pten for inducing and sustaining cell cycle arrest, cell polarity and lumen formation. Upon lrECM exposure in 3D, EpH4 cells give rise to quiescent polarized acini with a central lumen, recapitulating many cellular and molecular events observed during mammary gland morphogenesis *in vivo*^26^.

To better understand the role of Pten in the interplay between quiescence acquisition and maintenance, as well for establishing and sustaining apicobasal polarity and tissue architecture, we pharmacologically inhibited Pten in 3D epithelial acini in two distinct moments: during development and in mature structures (Fig. 7).

**Figure 7.**
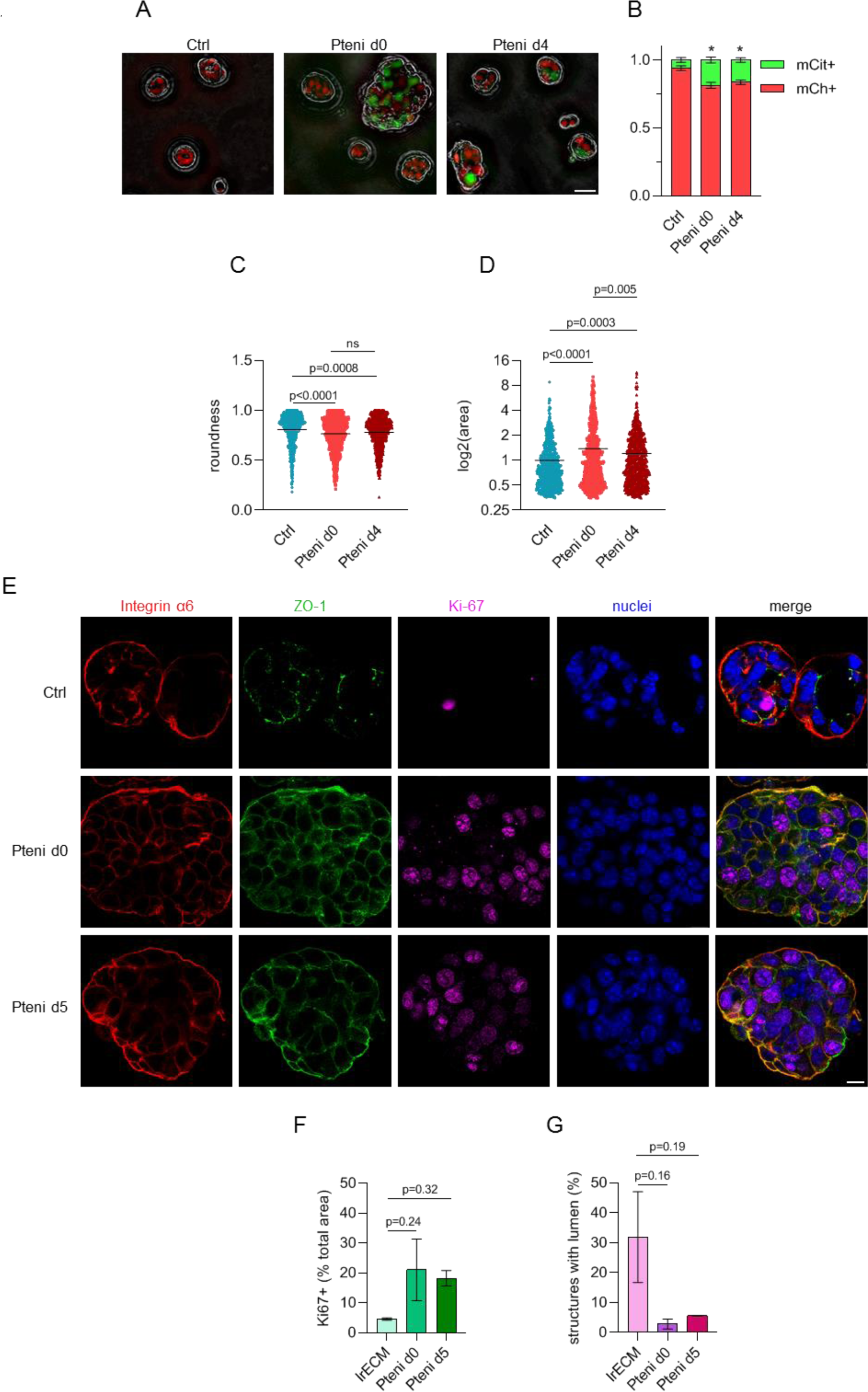
Pten inhibition affects quiescence and polarity in both developing and mature mammary 3D acini. A-D. EpH4-FUCCI cells were seeded on top of lrECM (matrigel) bedding and treated or not with 2.5 µM bpV(pic) (Pten inhibitor, Pteni) on the day of seeding (Pteni d0) or when the 3D structures were already established quiescent at day 4 (Pteni d4) and imaged at day 5 (n=2). A. Merged bright field and fluorescent images of representative structures for each treatment (Ctrl, Pteni d0 and Pteni d4). Red: Cdt1-mCherry FUCCI probe (cells in G1); green: mCitrine-geminin FUCCI probe (cells in S/G2/M). Scale bar 25 µm. B. Cell cycle quantification based on the expression of FUCCI probes (mean±SEM). C. Values of roundness of individual structures (each dot represents an individual 3D structure and the line marks the mean, total number of structures analysed per condition: Ctrl = 775, Pteni d0 = 812, Pteni d5 = 875). D. Normalized area (relative to the control). The values for area were plotted as log2(area), each dot represents an individual 3D structure and the line marks the mean, total number of structures analysed per condition: Ctrl = 775, Pteni d0 = 812, Pteni d5 = 875). E-G. EpH4 cell clusters grown in poly(HEMA)-treated plates were subjected to lrECM treatment (Ctrl), along with pharmacological inhibition of Pten with 2.5 µM bpV(pic) during acini morphogenesis (inhibitor added in the same day structures were treated with lrECM, Pteni d0) or when the structures were already mature (inhibitor added after 5 days of lrECM treatment, Pteni d5). Structures were collected and processed 6 days post lrECM treatment (n=2). E. Confocal images of 3D structures stained for integrin α6 (red, basal marker), ZO-1 (green, apical marker) and Ki-67 (magenta, proliferation marker). Scale bar 10µm. F. Percentage of Ki-67 positive signal relative to the total area in a given sample (mean±SEM). G. Percentage of structures with lumen (mean±SEM, total number of structures analysed per condition: Ctrl = 1747, Pteni d0 = 1610, Pteni d5 = 1482).

**Figure 8.**
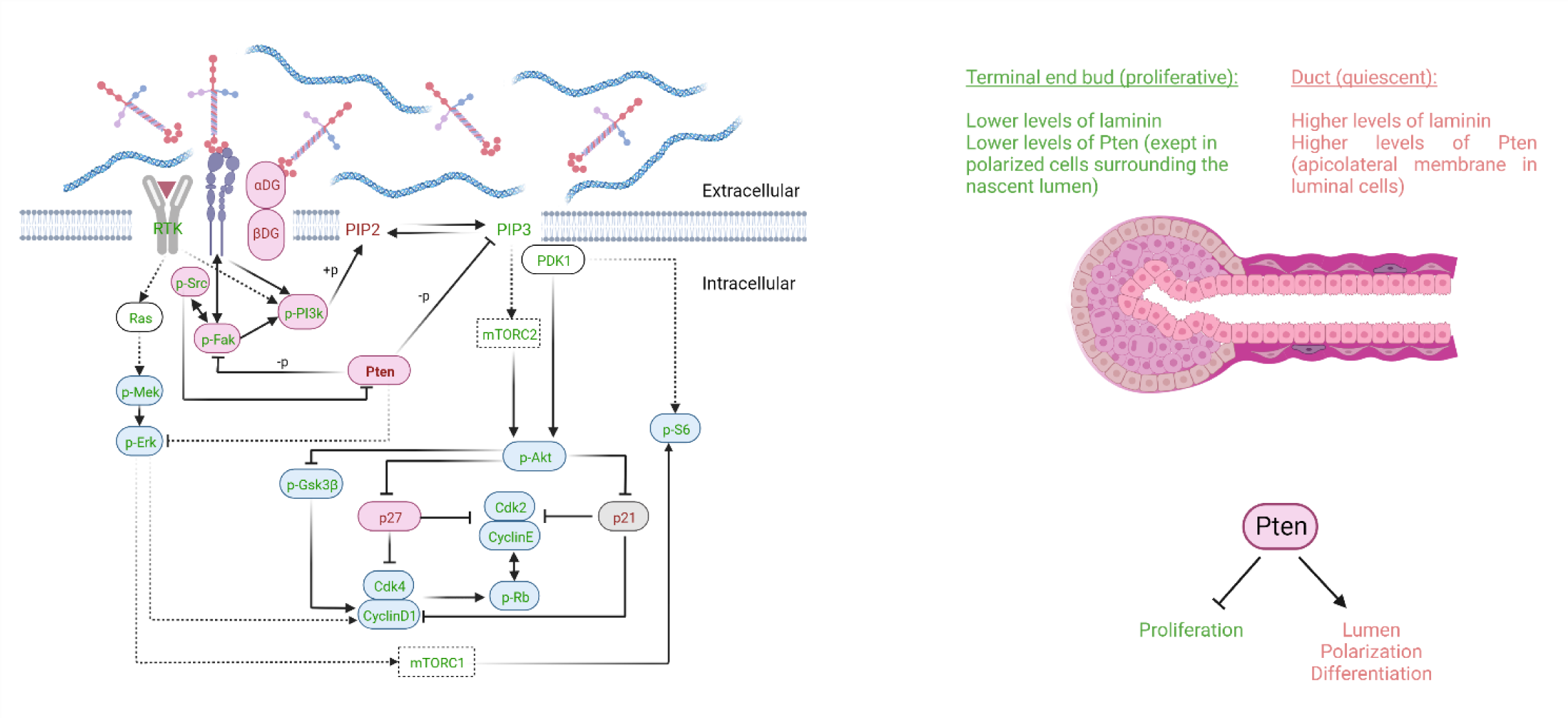
Proposed regulatory mechanism of modulation of mammary epithelial cell quiescence/proliferation decision by Pten. A. Molecular landscape of some cell cycle and proliferation regulators in laminin-induced quiescent mammary epithelial cells. Names of molecules classically associated with cell cycle progression are depicted in green, whereas molecules classically associated with cell cycle arrest are in red. Blue boxes: molecules that were found downregulated during lrECM-induced quiescence; red boxes: molecules that were found upregulated during lrECM-induced quiescence; gray boxes: molecules that exhibited to alteration during lrECM-induced quiescence; white boxes: not assessed; box with white/dashed line: protein complex, not assessed. B. Interplay between Pten and laminin-111 during mammary gland development to coordinate differentiation and cell cycle arrest.

In both cases, Pten inhibition led to increased proliferation, which was assessed via the expression of FUCCI sensors (Fig. 7 A-B) or by Ki-67 staining (Fig. 7 E-F). These data indicate that, in mammary epithelial cells, Pten activity is necessary to induce quiescence during development, and to sustain quiescence in mature structures (Fig. 7 A-B, E-F). Structures treated earlier (and therefore for longer) with Pten inhibitor were larger than those exposed later, although in both conditions treated structures were larger than control ones (Fig. 7 D).

EpH4 differentiated acini express apicobasal markers such as ZO-1 (apical) and integrin α6 (basal) (Fig. 7E, top panel), however, Pten inhibition during acini assembly led to the development of irregular structures which lacked a lumen and had widespread membrane staining for ZO-1 and integrin α6 (Fig. 7 A, C, E and G). Similarly, Pten inhibition in mature structures also disrupted apicobasal polarity and (Fig. 7 A, C-E and G). Interestingly, the effects of Pten inhibition on unleashing proliferation and disrupting tissue architecture were equally severe when inhibition occurred during development or in mature acini, (Fig. 7 A-C, E and G), which is consistent with a role of Pten in inducing and sustaining quiescence and tissue architecture.

## Discussion

By using a relevant cell culture model, where even in the presence of growth factors, exposure to a basement membrane surrogate^59^ triggers quiescence and differentiation in mammary epithelial cells, akin to what occurs *in vivo*^1,11,21,57^. We initially mapped mRNA, protein and phosphorylation levels of several cell cycle regulators in quiescent cells. The diversity of alterations found in this molecular landscape helps to explain why overexpressing a CKI or downregulating cyclins/Cdks might lead to irreversible cell cycle arrest, not fully recapitulating the transient state of quiescence^60,61^.

By combining the snapshot of the status of cell cycle regulators in lrECM-induced quiescent cells, the use of fluorescent cell cycle indicators, perturbation with specific inhibitors and the pertinent literature, we can infer signalling networks that might lay the molecular foundation for specific traits of quiescent cells. For example, in order to proliferate, cells need to grow throughout G1 until they achieve a specific targeted size^62–64^. Thus, although cell growth and cell proliferation are separable events, there is high coupling degree between cell size and cell cycle progression^65^. Mechanisms underlying cell size control are poorly understood^64^, but mTOR is a clear positive regulator of cell growth^43^. The reduced size observed in quiescent cells, along with our findings of reduced mTOR signalling upon lrECM treatment and induction of cell cycle arrest by mTORC1/mTORC1/2 inhibitors, combined with previous findings^43,65^ point out that reduced cell growth might be one of the mechanisms behind cell cycle arrest in lrECM-induced quiescence in mammary epithelial cells.

Another intriguing observation is the differential effects of CDK2 and CDK4 inhibition in cell proliferation and morphology, together with the interplay between cyclins, CDKs and CKIs during lrECM-induced quiescence. A few things need to be considered in order to better explore these data. First, in the classical model of cell cycle progression, in response to mitogenic stimuli during G1, cyclinD-CDK4 complexes phosphorylate and inactivate Rb, which releases E2F transcription factors, which in turn, switch on the expression of a range of positive regulators of the cell cycle, including cyclin E, which in complex with CDK2 further phosphorylates and inhibits Rb^31,66^. From this point on, known as the “restriction point” and molecularly characterized by Rb hyperphosphorylation, cells are committed to proliferation and will progress through cell cycle regardless of mitogenic stimuli^66,67^. However, it has been recently demonstrated that molecular events occurring much earlier than Rb hyperphosphorylation, including the degree of CDK2 activity in newly born cells, are the determinant factors behind cell cycle commitment^23^.

Second, although the CKIs p21 and p27 supress CDK2 activity, they are required for the assembly of cyclin D-CDK4 complexes. In this context, p21 is generally an inhibitor or a poor activator, while p27, depending on its phosphorylation status, can be either an inhibitor or a strong activator for cyclin D-CDK4 complexes^49,68^. Thus, in addition to its kinase activity, cyclin D-CDK4-p27 complexes would contribute for cell cycle progression by titrating p27 away from CDK2 complexes^49,68^.

Although a much more in-depth analysis of these mechanisms would be necessary to draw any conclusions, it is tempting to speculate that during lrECM-induced cell cycle arrest, inhibition of CDK2 activity could be a core event for acquiring and sustaining the quiescent phenotype. LrECM-induced inhibition of CDK2 activity, due to both reduced levels of cyclin E and CDK2, along with increased p27, in a context where cyclin D1 and CDK4 were also reduced, would allow for cell cycle arrest even in the presence of mitogens. Which of these events comes first, how this is regulated at the molecular level and whether this stands true for lrECM-induced quiescence would need extensive investigation and is beyond the scope of the present work.

The apparent discrepancy between the activation status of some upstream (Fak, Src and PI3k) and downstream (Akt, mTOR and Mapk) regulators of cell cycle progression could also be consistent with the idea that proliferative signals from the ECM are sensed by quiescent cells, but that proliferative signalling effectors are actively shut down by laminin-triggered alterations.

Depending on spatiotemporal regulation, signalling integration and cellular sub localization, Fak, Src and PI3k are involved in a range of cellular processes other than proliferation, such as cell survival^69^, invasion^70^, migration^71^, polarity^17^ and lumen formation^44^. Hence, shutting down the proliferative signalling arms of these pathways might open space to the rise of other signalling routes, and for instance, shifting cell phenotype towards differentiation.

Our findings point out that Pten might, at least in part, fulfil the role of shifting proliferative signalling towards a differentiation program. Pten was upregulated at the protein level in quiescent cells (Fig. 2), its inhibition led to increased proliferation, restored Akt/mTORC1/2 and Mapk signalling in the presence of quiescence-inducing signals from lrECM in mammary epithelial cells and disrupted quiescence, polarity and tissue architecture in 3D acini (Fig. 3,4 and 7).

Pten is a well known tumour suppressor, and its role in restraining cell proliferation has been extensively demonstrated in a wide range of tissues and cell types^14^. The PI3k pathway is one of the most mutated pathways in cancer, with Pten figuring, in fact, as the most frequently altered PI3K pathway member^72^. Pten loss has been associated with increased proliferation due to upregulation of PI3k/Akt/mTORC1/2 and MAPK signalling pathways^28,73^, but even discrete alterations in Pten levels can trigger dramatic changes and cell behaviour and increase the risk of developing cancer^14^. Src has been reported as a negative regulator of Pten activity, thus, one of the mechanisms by which Src inhibition might lead to cell cycle arrest is by increased Pten activity and consequent repression of the PI3K/mTOR/Akt axis^74^. Interestingly, mTORC1 hyperactivity downstream of Pten loss has been shown to trigger a feedback loop that restrains full PI3K activation^75^. The existence of this feedback loop helps to explain the lack of further phosphorylation of Akt^T308^ in control cells, and might indicate that this feedback loop could be repressed upon lrECM-treatment, where Pten inhibition led to increase in phospho-Akt^T308^ (Fig. 3). We can not rule out that a similar negative feedback loop, restraining activation in the absence of quiescent cues but inhibited by lrECM, might exist for Mapk, once Pten inhibition was able to increase the levels of phospho-Erk in lrECM-treated cells but not in control cells (Fig. 3). Confirming these hypotheses would require additional investigation, but such negative feedback loops could be important for allowing cells to grow and get to their target size prior to cell division, or for the occurrence of other cell behaviours, including migration and invasion.

The mechanisms by which laminin upregulates Pten and if this regulation follows the rules of dynamic reciprocity^4^ remain unclear, but, in 3D culture of mammary epithelial cells, Pten levels have been linked to increased dystroglycan receptor expression^76^, as well as to e-cadherin expression and localization^77^. Tissue architecture integrity and the levels and localization of Pten have been pointed out to exist in dynamic reciprocity^77^. We did find increased levels of dystroglycan receptor upon lrECM-induced quiescence (Fig. 2A, D and E). The dystroglycan receptor has two subunits encoded by a single transcript^78^. The extracellular subunit α-dystroglycan is subjected to posttranslational modifications through glycosylation, which confers its ability to bind laminin^5,6^. The intracellular subunit β-dystroglycan usually binds to α-dystroglycan and to cytoskeleton proteins, although it can also be found in the nucleus^5,79^. Dystroglycan receptors is associated to attenuation of integrin signalling^80^, cell cycle arrest^30^ and polarity^76^ in mammary epithelial cells treated with laminin-rich ECM. Therefore, increased expression of Pten and dystroglycan receptor and might be cooperating to trigger and sustain quiescence in growth suppressive microenvironments.

Correlations between BM thickness and some regulators of proliferation and migration, including nuclear actin^10^ and nuclear galectin-1^12^, have been reported in the developing mammary gland, and our data indicate that Pten might also fall into this category. Endogenous Pten displayed differential levels and staining patterns in TEBs versus ducts of the developing mammary gland *in vivo*, which correlated to the degree of laminin deposition in these structures (Fig. 5). Interestingly, cells surrounding the nascent lumen in TEBs exhibited apicolateral Pten staining, the same pattern found in differentiated, luminal cells in the ducts (Fig. 5 and 6). These data might indicate that, in TEBs, cells surrounding the lumen already show some degree of polarization and maybe even differentiation, whereas cells in other areas do not. Indeed it has been shown that even though the TEB is a highly proliferative structure within the developing mammary epithelia, proliferation was not observed in luminal epithelial cells surrounding the nascent lumen^81^, and our data indicate that Pten might have a role in this phenomenon.

Pten in the apicolateral membrane, either in ducts or in cells surrounding the lumen in TEBs, strongly colocalized with the actin belt (Fig. 6), resembling the staining of other classical apicolateral markers of polarized cells, such as ZO-1 and e-cadherin^82,83^. Whether cell differentiation precedes Pten apicolateral polarization, or if it is the other way around, remains to be experimentally demonstrated, however, Pten most likely acts on the onset of polarity acquisition rather than being a late factor recruited to mature junctional complexes.

It has been shown that Par3, which has a core role in the establishment of cell polarity, and e-cadherin, a key regulator of epithelial tissue integrity, are both able to recruit Pten to cell-cell junctions, which is required for cell differentiation^84,85^. Polarization of Pten to the apicolateral membrane might contribute for the establishment of lipid domains^86,87^ and for the compartmentalization of PI3k activity in quiescent, differentiated cells^85^. In fact, PIP2, the lipid product of Pten activity, has been proposed to be an apical marker itself^15,87^. PIP2 is known to recruit to the membrane a range of proteins with relevance for cell polarization, tissue architecture and cell cycle arrest, including annexin 2 and Cdc42^15^. Additionally, even though PI3k inhibition leads to phenotypic reversion of malignant T4-2 breast cells, some asymmetric PI3k activity seems necessary for polarity, once Pi3k and its product PIP3 localize to the basal surface of S1 and reverted T4-2 acini^17^. Indeed, EpH4 acini also display active PI3k polarized to the basolateral membrane, and activation of the PI3k-Rac1 axis is required for complete differentiation and milk protein synthesis^21,26^, thus, polarized activity of PI3k to the basal surface might trigger survival and differentiation, rather than proliferation.

Our data support a role for Pten not only in proliferation inhibition, but also in lumen assembly and maintenance in the developing mammary gland, and are consistent with previous findings^15,16^. Apicolateral Pten was reported in several 3D polarized epithelial structures such as MDCK (kidney) and T84 (colon) cysts^15^, chick epiblast^88^, as well as in human and murine mammary epithelial cells^16,77^, but, to our knowledge, for the first time in the developing mammary gland *in vivo*.

Facing these results, an old, although not fully understood question came to mind: what comes first: polarity or cell cycle arrest? And at which point does Pten come to play?

Cell polarity and cell cycle arrest are interrelated, PI3k-modulated, but molecularly distinct events, where inhibition of the PI3k downstream effectors Akt and Rac1 were linked to proliferation arrest and polarity, respectively^17^. Similarly, by using 3D acini, Leslie and collaborators have shown that Pten requires both lipid and protein phosphatase activity to control lumen formation through a mechanism that does not correlate with its ability to control AKT. Interestingly, in NMuMG acini, despite of displaying increased cell proliferation Pten knockdown or overexpression of p110a/PIK3CA oncogenic mutants disrupt lumen formation, whereas a constitutively activated Akt does not^16^. These data further support the notion that different pathways downstream PI3k govern the regulation of lumen formation and proliferation, and that Pten might be a key regulator of both. In this sense, our data in 3D acini models provide further evidence that Pten simultaneously coordinates proliferation and normal tissue architecture during acini development, and sustains quiescence and tissue architecture in mature, quiescent acini (Fig. 7).

The data we presented here add up to the evidence that a laminin-rich ECM, found in the basement membrane, induces quiescence and differentiation in mammary epithelial cells by differentially modulating the expression and activity of cell cycle regulators at the transcription and protein level. This panorama, along with the extensive previous work done by many others, provide some hints on molecules, pathways and mechanisms involved in microenvironmental control of quiescence in higher organisms, which could be explored in studies to come.

In the context of epithelial cell quiescence and differentiation, Pten seems to be one of the key molecular players. Pten levels and localization pattern were correlated to basement membrane deposition and quiescence acquisition *in vivo*, where Pten seemed to induce quiescence, polarity and lumen formation during mammary gland development, and sustain this quiescent and differentiated phenotype in mature mammary epithelial structures.

Lastly, the notion that lrECM-triggered quiescence involves several pathways and has many layers of regulation provides an explanation for the robustness of this system in suppressing epithelial cell proliferation, which could ultimately be linked to cancer incidence being lower than expected^89^ and even for the need of more than one perturbation to bypass microenvironmentally-induced suppression of cancer development^90^. In this sense, disruptions in master regulators such as Pten, which regulate several signalling pathways and phenotypical features required for tissue homeostasis and tumour suppression, could indeed be drivers of malignant transformation.

## Materials and methods

### Reagents

Primary and second antibodies used in this work are listed on Table 1, and Table 2, respectively. Table 3 brings information on the pharmacological inhibitors used. Additional reagents used are listed on Table 4 and plasmids are listed on Table 6.

**Table 1.** List of primary antibodies

| Target | Host | Manufacturer | Catalog number | Working concentration & application |
| --- | --- | --- | --- | --- |
| α-Tubulin | Mouse | Santa Cruz | sc-5286 | 1:5000 (WB) |
| Lamin B1 | Goat | Santa Cruz | sc-6217 | 1:1000 (WB) |
| α-dystroglycan (VIA41) | Mouse | DSHB - Iowa | | undiluted (WB) |
| α-dystroglycan (IIH6) | Mouse | DSHB - Iowa | | undiluted (WB) |
| β-dystroglycan | Mouse | Biolegend | 850202 | 1:1000 (WB) |
| p-Fak<sup>Y397</sup> | Rabbit | Life Technologies | 44-624G | 1:500 (WB) |
| p-Fak<sup>Y861</sup> | Rabbit | Life Technologies | 44-626G | 1:500 (WB) |
| Fak | Rabbit | Cell Signalling | 3285S | 1:500 (WB) |
| p-Src<sup>Y416</sup> | Rabbit | Cell Signalling | 2101 | 1:500 (WB) |
| Src | Rabbit | Cell Signalling | 2109 | 1:1000 (WB) |
| p-PI3K(p85)<sup>T467</sup>/(p55)<sup>T199</sup> | Rabbit | Cell Signalling | 4228S | 1:500 (WB) |
| PI3K(p85) | Rabbit | Cell Signalling | 4292S | 1:1000 (WB) |
| Pten | Rabbit | Cell Signalling | 9559S | 1:500 (WB), 1:50(IF) |
| p-Akt<sup>T308</sup> | Rabbit | Cell Signalling | 4056S | 1:500 (WB) |
| p-Akt<sup>S473</sup> | Rabbit | Cell Signalling | 4060 | 1:500 (WB) |
| Akt | Rabbit | Cell Signalling | 9272 | 1:1000 (WB) |
| p-S6<sup>S240/244</sup> | Rabbit | Cell Signalling | 5364 | 1:1000 (WB) |
| S6 | Rabbit | Cell Signalling | 2217L | 1:1000 (WB) |
| p-GSK3B<sup>S9</sup> | Goat | Santa Cruz | 11757 | 1:500 (WB) |
| GSK3B | Mouse | Santa Cruz | 377213 | 1:1000 (WB) |
| p-MEK1/2<sup>S217/221</sup> | Rabbit | Cell Signalling | 9154 | 1:500 (WB) |
| MEK1/2 | Rabbit | Cell Signalling | 9126 | 1:500 (WB) |
| p-ERK<sup>Y202/204</sup> | Rabbit | Cell Signalling | 9101 | 1:1000 (WB) |
| ERK | Rabbit | Cell Signalling | 9102 | 1:1000 (WB) |
| CyclinD1 | Rabbit | Thermo Fisher | MA5-16356 | 1:1000 (WB) |
| CDK4 | Rabbit | Cell Signalling | 12790 | 1:250 (WB) |
| CyclinE1 | Rabbit | Santa Cruz | 481 | 1:250 (WB) |
| CDK2 | Rabbit | Santa Cruz | 163 | 1:250 (WB) |
| p27 | Rabbit | Abcam | ab32034 | 1:500 (WB) |
| p21 | Rabbit | Abcam | ab7960 | 1:250 (WB) |
| pRb<sup>S807/811</sup> | Rabbit | Cell Signalling | 8516 | 1:1000 (WB) |
| ZO-1 | Mouse | Thermo Fisher | 61-7300 | 1:50(IF) |
| α6 Integrin | Rat | BD Biosciences | 555734 | 1:50(IF) |
| Ki-67 | Rabbit | Abcam | Ab16667 | 1:50 (IF) |
| Laminin 111 | Goat | Bissell's lab |  | 1:500 (IF) |
| Rabbit IgG |  | Cell Signalling | 3900 | 1:100 (IF) |

**Table 2.** List of secondary antibodies

| Target | Conjugation | Manufacturer | Working concentration & application |
| --- | --- | --- | --- |
| mouse IgG light chain | HRP | Jackson Labs | 1:20000 (WB) |
| rabbit IgG light chain | HRP | Jackson Labs | 1:20000 (WB) |
| goat IgG light chain | HRP | Jackson Labs | 1:20000 (WB) |
| rabbit IgG | AlexaFluor 488 | Thermo Fisher | 1:400 (IF) |
| rat IgG | AlexaFluor 546 | Thermo Fisher | 1:400 (IF) |
| Mouse IgG | AlexaFluor 647 | Thermo Fisher | 1:400 (IF) |

**Table 3.** List of specific pharmacological inhibitors

| Inhibitor | Target | Manufacturer | Catalog number | Working concentration |
| --- | --- | --- | --- | --- |
| FAK inhibitor 14 | Fak | Sigma-Aldrich | SML0837 | 1 μM |
| PP2 | Src | Calbiochem | 172889-27-9 | 2 μM |
| CPD22 | ILK | Sigma-Aldrich | 407331 | 0.25 μM |
| Tofacitinib | Jak/Stat | Selleckchem | CP-690550 | 20 μM |
| LY294002 | PI3K | Invitrogen | PHZ1144 | 20 μM |
| bpV(pic) | Pten | Sigma-Aldrich | SML0885 | 2.5 μM |
| Rapamycin | mTORC1 | LC Laboratories | R-5000 | 0.2 μM |
| Torin 2 | mTORC1/2 | TOCRIS | S2817 | 0.4 μM |
| MK 2206 | Akt | Selleckchem | S1078 | 2 μM |
| PD98059 | Mek | Invitrogen | PDZ1164 | 20 μM |
| Palbociclib | Cdk4 | Sigma-Aldrich | PZ0383 | 1 μM |
| CDK2 inhibitor III | Cdk2 | Calbiochem | 238803 | 60 μM |

**Table 4.** Additional reagents and materials

| Reagent/material | Manufacturer | Catalog number |
| --- | --- | --- |
| DAPI | Sigma-Aldrich | D9542 |
| AlexaFluor 647 Phalloidin | Invitrogen | A22287 |
| Hoechst 33342 | Invitrogen |  |
| Insulin | Sigma-Aldrich | I0516 |
| Gentamycin | Gibco | 15710072 |
| RNAse A | Sigma-Aldrich | 10109142001 |
| Propidium iodide (PI) | Invitrogen | P4170 |
| ProLong™ Diamond Antifade Mountant | ThermoFisher | P36970 |
| DMEM/F12, phenol red | Gibco | 12-500-062 |
| DMEM/F12, no phenol red | Gibco | 21041025 |
| Hydrocortisone | Sigma-Aldrich | H0888 |
| Trypsin-EDTA 0.05% | Gibco | 25300054 |
| Corning® Matrigel® Matrix, Growth Factor Reduced | Corning | 356231 |
| Polybrene | Sigma-Aldrich | H9268 |
| Polyethylenimine (PEI) | Polysciences | 23966 |
| Tissue Tek O.C.T | Sakura | 4583 |
| Starfrost Microscope Slides | Knittel Glaser |  |
| Odyssey Blocking buffer | LI-COR | 927-50003 |
| RNAeasy kit | Qiagen | 74106 |
| Ambion Turbo DNase kit | Invitrogen | AM1907 |
| Superscript II Reverse Transcriptase | ThermoFisher | 18064014 |
| OligoDT primers | ThermoFisher | 18-418-020 |
| Random hexamer primers | ThermoFisher | N8080127 |
| Power SYBR™ Green PCR Master Mix | Applied Biosystems | 4368702 |
| SuperSignal™ West Dura Extended Duration Substrate | Life Technologies | 34076 |
| DC Protein Assay kit | BioRad | 500-0116 |
| Precision Plus Protein™ Dual Color Standards | BioRad | 1610374 |

### Cell culture

Murine mammary epithelial cells EpH4, EpH4-ES-FUCCI and EpH4-DHB-mVen-mCh-CDK4.KTR-H2B-mTurq cells were routinely grown in DMEM/F12, 2% FBS, 5µg/mL insulin, 50µg/mL gentamycin at 37°C and 5% CO_2_ in a humidified incubator. Cells were passaged when necessary (every 2-3 days) using Trypsin 0.05%-EDTA.

#### Laminin-rich ECM overlay differentiation assay

EpH4 cell quiescence induction and lactogenesis differentiation upon overlay treatment with laminin-rich ECM has been described elsewhere^10^. Briefly, cells were seeded at 2000 cells/cm^2^ (unless otherwise specified). In the following day, media was changed for GIH media (DMEM/F12, 50 µg/mL gentamycin, 5 µg/mL insulin, 1 µg/mL hydrocortisone). 24h later, cells were treated or not with 2% growth factor reduced matrigel (lrECM) and/or 3 µg/mL prolactin (Plr and lrECM+Plr), and, when appropriate with specific inhibitors. Cells were harvested or imaged 24-48h post treatment (depending on the assay).

### Cell cycle analysis via PI staining and flow cytometry

EpH4 cells subjected or not to lrECM overlay treatment for 48h. Cells were then detached and pelleted by centrifugation, washed with cold PBS, fixed in 70% cold ethanol and stored for 24h at −20°C. Fixed cells were incubated with RNase A (0.2 mg/mL) at 37°C for 30 min and stained with Propidium iodide (PI) (1 μg/mL). DNA content of cells (20,000 events) was analysed using an ACCURI C6 flow cytometer. Data were plotted on GraphPad Prism and statistical differences between treatments for each cell cycle phase were assessed by Two-way ANOVA followed by Sidak’s multiple comparisons test. p<0.05 was considered significant.

### Cell density and live and dead assay

EpH4 cells subjected or not to lrECM overlay treatment for 48h. Cells were then stained with the DNA-binding dyes PI (5 μg/mL) and Hoechst 33342 (5 μg/mL) for 15 min. Cells were then imaged on a customized TissueFAXS i-Fluo system (TissueGnostics) mounted on a Zeiss AxioObserver 7 microscope (Zeiss), using 20× Plan-Neofluar (NA 0.5) objective and an ORCA Flash 4.0 v3 camera (Hamamatsu). 8X8 adjacent microscope fields were acquired per well using automated autofocus and image acquisition settings. Thousands of cells per sample were analysed using a customized pipeline on StrataQuest software (TissueGnostics), which can be made available upon request. Details on parameters used for nuclei segmentation and detection on StrataQuest can be found at Russo *et al*., 2021^91^. Individual nuclei for all cells were detected in the Hoechst channel, which was used to build a nuclear mask where the Propidium iodide signal (dead cells) was detected. The relative cell density (number of cells based on the Hoescht signal/total area imaged) was calculated for control (normalized to 1) or lrECM-treated cells. Data were plotted on GraphPad Prism and statistical differences between treatments were assessed by Student’s t-test. p<0.05 was considered significant. The percentage of dead cells was calculated as Propidium iodide positive cells/all cells (Hoechst). Data were plotted on GraphPad Prism and statistical differences between treatments for the percentage of live or dead cells were assessed by Two-way ANOVA followed by Sidak’s multiple comparisons test. p<0.05 was considered significant. These experiments were performed in triplicate wells in three biological replicates.

### FUCCI-based cell cycle analysis

For these experiments, EpH4 cells constitutively expressing the fluorescence cell cycle indicator FUCCI^92^ were used. EpH4-ES-FUCCI cells were generated by transfecting EpH4 cells with the ES-FUCCI plasmid (Table 6), followed by hygromycin B selection (500µg/mL, 10 days) and aseptic cell sorting of fluorescent cells.

EpH4-ES-FUCCI cells were seeded into 96 well plates and subjected to the lrECM overlay assay (control and lrECM). 48h post treatment, cells were incubated with Hoechst (5 μg/mL) for 15 min and imaged on a Leica DMi8 wide-field fluorescence microscope (e-signal lab/USP) using the LasX Navigator Application (Leica Microsystems). The DMi8 microscope was coupled with an incubation system which allowed live cell imaging at 37°C and 5% CO_2_. Each well was imaged in 3x3 adjacent microscope fields, using the 10x objective and specific fluorescence filters (DAPI for Hoescht; Txr for mCherry and YFP for mCitrine). Images were analysed on StrataQuest software (TissueGnostics) using a customized pipeline, which can be provided upon request. Details on parameters used for nuclei segmentation and detection on StrataQuest can be found at Russo *et al*., 2021^91^. In brief, nuclei were detected in the Hoechst channel and a mask was created, allowing the measurement of the mean fluorescent intensity signal of the FUCCI probes (Cdt1-mCherry and Geminin-mCitrine) for thousands of cells per sample. The percentage of mCherry+ (cells in G1), mCitrine+ or double-positive (cells in S/G2/M) or double negative cells (cells that had just finished M) was determined for each well. Data were plotted on GraphPad Prism and statistical differences between treatments for each cell cycle phase were assessed by Two-way ANOVA followed by Sidak’s multiple comparisons test. p<0.05 was considered significant. These experiments were performed in triplicate wells in three biological replicates.

### CDK2 and CDK4 activity

CDK2 and CDK4 activities were evaluated using kinase translocation reporters (KTR), as previously described^23,24^. EpH4-DHB-mVen-mCh-CDK4.KTR-H2B-mTurq cells were generated by simultaneous infection of EpH4 cells with lentiviruses containing either DHB-mVen-mCh-CDK4.KTR or the H2B-mTurq constructs in the presence of 8 µg/mL Polybrene. Lentiviruses were previously packaged by HEK293 FT cells transfected with polyethylenimine (PEI), the packaging plasmids psPAX2 and pMD2.G and pLenti-DHB-mVenus-p2a-mCherry-CDK4KTR or CSII-EF1-H2B-mTurquoise (Table 6), following instructions from Addgene. Cells were then subjected to aseptic sorting for selecting double-infected cells (expressing mVenus and mCherry, as well as mTurquoise).

EpH4-DHB-mVen-mCh-CDK4.KTR-H2B-mTurq cells were plated into 96 well plates and subjected to lrECM overlay assay (control and lrECM). 48h post treatment, cells were imaged on a Leica DMi8 wide-field fluorescence microscope (e-signal lab/USP) using the LasX Navigator Application (Leica Microsystems). Each well was imaged in 3x3 adjacent fields of view, using the 10x objective and specific fluorescence filters (CFP for mTurquoise; Txr for mCherry and YFP for mVenus). CDK2 and CDK4 activities were calculated based on the guidelines from Spencer *et al.*, 2013^23^ and Yang *et al*., 2020^24^, the developers of these specific probes. Images were analysed on StrataQuest software (TissueGnostics) using a customized pipeline, also available upon request. Details on parameters used for nuclei segmentation and detection on StrataQuest can be found at Russo *et al*., 2021^91^. In brief, nuclei were detected in the mTurq channel and a nuclear mask was created. In order to measure the cytoplasmic intensity of CDK2 and CDK4 probes, another mask, corresponding to a cytoplasmic ring was created (2 µm around the nucleus or less if touching another cytoplasmic ring or nuclei, starting 0.5 µm away from the nucleus). The mean intensity of fluorescent signals (mVenus for CDK2 and mCherry for CDK4) were obtained for the nucleus and for the cytoplasmic ring for thousands of individual cells per sample. For each cell, CDK2 activity was calculated as the ratio of the mean fluorescent intensity of mVenus in the cytoplasmic ring and in the nucleus. Once CDK2 can at some extent phosphorylate the CDK4 probe^24^, CDK4 activity was calculated as the cytoplasmic/nuclear ratio of the mean fluorescent intensity of mCherry, minus the CDK2 activity in that cell multiplied by a correction factor that was previously calculated according to Yang *et al*., 2020^24^ (CDK4 activity = mCherry cyto/mCherry nuc – CDK2 activity*0.05). Data were plotted as a “Superplot”^93^ on GraphPad Prism and statistical differences between treatments for the activity of each CDK in control or treated cells were assessed by One way ANOVA followed by Sidak’s multiple comparisons test. p<0.05 was considered significant. These experiments were performed in triplicate wells in three biological replicates.

### Inference of cell size via flow cytometry and microscopy

EpH4 cells were subjected to lrECM overlay assay for 48h and cell size was estimated according to the forward scatter area (FSC-A). Once freely cycling cells (control) would have significantly more cells in S/G2/M (which are larger in size) than quiescent (lrECM-treated) cells, in order to do a fair comparison, both control and lrECM-treated cells were gated in G1 according to DNA content (PI staining).

Similarly, EpH4-FUCCI cells were subjected to lrECM overlay assay for 48h, counterstained with Hoechst (5 µg/mL, 15 min) and imaged (as per described in the “FUCCI-based cell cycle analysis” section). Cell size was estimated according to the area of the nucleus. Images were analysed on StrataQuest software (TissueGnostics), and once again, only cells in G1 (mCh+) were considered for this analysis.

Data were plotted as the mean size (normalized to control) on GraphPad Prism and statistical differences between treatments were assessed by Student’s t-test. p<0.05 was considered significant. These experiments were performed in three biological replicates.

### Gene expression analysis

EpH4 cells were subjected to lrECM overlay and/or prolactin treatment for 48h. Cells were then harvested for RNA extraction, reverse transcription and RT-qPCR. This experiment has three biological replicates.

Gene expression was assessed by RT-qPCR, following the MIQE guidelines^94^. Briefly, total RNA was extracted using RNeasy kit, quantified and treated with Turbo DNAse kit according to the manufacturer’s instructions. DNAse-treated RNA was reserve transcribed into cDNA using the Superscript II Reverse Transcriptase, oligoDT and random hexamer primers (Thermo Scientific), following the manufacturer’s guidelines.

RT-qPCR was performed in triplicates using 5 ng of cDNA, Power SYBR™ Green PCR Master Mix and specific primers (listed in table 5) into a 10 µL reaction. The reactions were ran in the AB-7500 real-time thermocycler (Applied Biosystems) using standard settings. B2m was used as the endogenous control and mRNA fold-changes (FC) were calculated according to the Pfaffl method^95^.

**Table 5.**
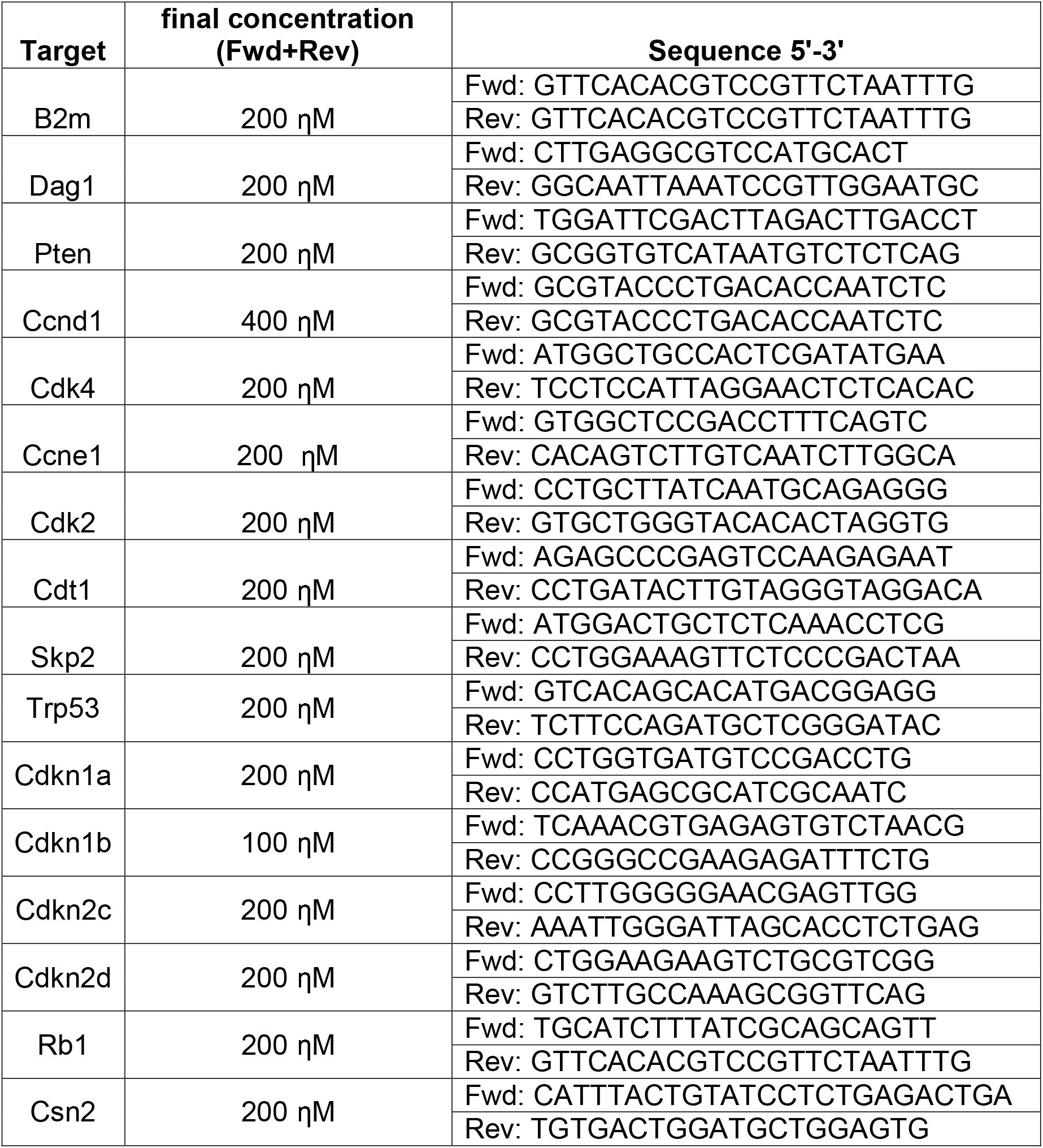
Primers used for RT-qPCR

**Table 6.**
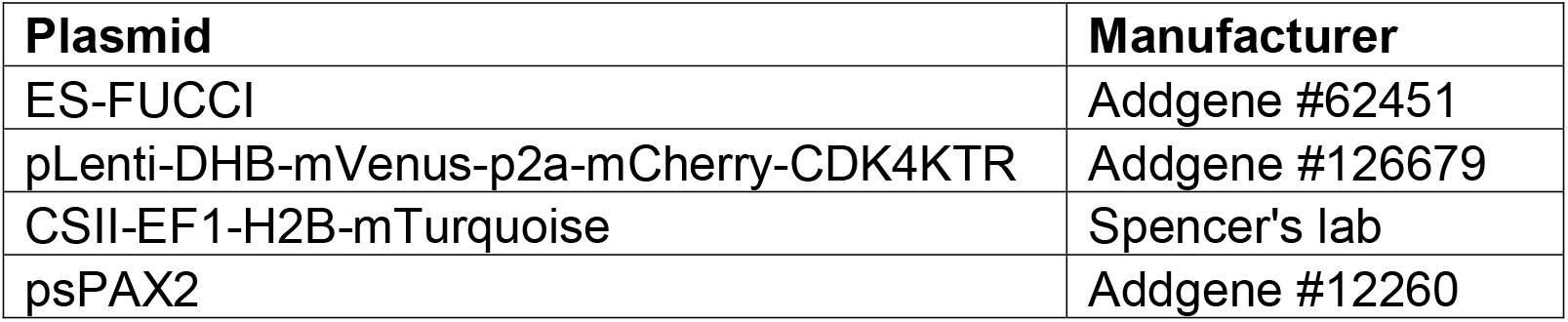
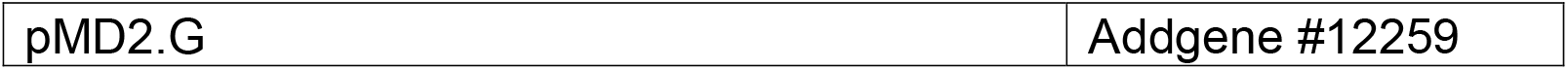
Plasmids

| Plasmid | Manufacturer |
| --- | --- |
| ES-FUCCI | Addgene #62451 |
| pLenti-DHB-mVenus-p2a-mCherry-CDK4KTR | Addgene #126679 |
| CSII-EF1-H2B-mTurquoise | Spencer's lab |
| psPAX2 | Addgene #12260 |

Statistical analysis was carried out on Graphpad Prism using one way ANOVA, followed by Dunnett’s multiple comparison test (comparing all samples to the Control, p<0.05 was considered significant). Data were presented as log2FC and –log10pValue using the Online Software Morpheus (Broad Institute)^96^.

### Protein expression analysis

EpH4 cells were subjected to lrECM overlay and/or prolactin treatment for 48h. Cells were then lysed with boiling 1x Laemmli Buffer (62.5 µM Tris, 2% SDS, 10% glycerol) and the obtained lysate was further incubated at 100°C for 15 min and stored at −20°C. Protein was quantified using the DC Protein Assay kit (BioRad) according to the manufacturer’s instructions.

The levels of specific proteins were then assessed by western blotting. For loading the gel, 1 µL of 0.15% bromophenol blue in 2-Mercaptoethanol was added to 20 µg of protein in 1 x Laemmli Buffer for a total volume of 30 µL. Samples were boiled at 100°C for 10 min and ran in into Tris-glycine polyacrylamide gels (8%, 10%, 12% or 15%, depending on the molecular weight of the protein of interest). Resolved proteins were transferred overnight into a PVDF membrane (0.22µm, Millipore) followed by 30 min incubation in blocking buffer (TBS, 0.2% Tween-20 with 5% BSA). Membranes were incubated overnight in blocking buffer containing primary antibodies (listed in Table 1), followed by incubation with the appropriate HRP-conjugated second antibody (listed in Table 2) for 1h at room temperature. HRP was detected by SuperSignal™ West Dura Extended Duration Substrate (Thermo-Fisher Scientific) and the chemiluminescence signal was captured with a ChemiDoc MP Imaging System (BioRad).

When necessary, membranes were subjected to a stripping protocol to remove antibodies and allow the reprobing with antibodies against another protein (this was used particularly to assess the total levels of a given protein after assessing its phosphorylated levels). Stripping consisted in 2x 10 min in distilled water, followed by 1 min in stripping buffer (25 mM Glycine, 1% SDS, pH 2) and 4 x 5 min in TBS, 0.2% Tween-20. Membranes were then blocked for 30 min and subjected to standard primary/HRP-secondary antibody incubation and detection.

For the relative quantification of protein levels, the optical density of each band was determined with ImageJ. The endogenous control (lamin B or α-tubulin, depending on the experiment) was also quantified and the relative expression was determined as the ratio between target protein level and the endogenous control. For phosphorylated proteins, we used the ratio phosphorylated protein/total protein/endogenous control. Phospho-Rb and phospho-Pi3k(p55) were normalized only against the endogenous control once we could not assess the total levels of these proteins.

Data were analysed on Graphpad Prism. Fold change differences of a given protein between two groups were evaluated by unpaired t-test (p<0.05 was considered significant). The obtained data were compiled and displayed in Volcano Plots (log2FC *vs* –log10pVal). Comparisons between multiple groups were performed by one way ANOVA, followed by Tukey post-test (p<0.05 was considered significant). This experiment has at least three biological replicates for each protein and condition analysed.

### Treatment with inhibitors and live cell imaging for FUCCI-based cell cycle analysis

EpH4-ES-FUCCI cells were seeded into 96 well plates (Corning) at a density of 2000cells/well. In the following day, media was changed to GIH without phenol red. 24h later, cells were treated or not with 2% Matrigel and/or specific inhibitors (Table 3). 24h post treatment, cells were incubated with Hoechst, imaged and analysed as per described in the “FUCCI-based cell cycle analysis” section. The relative cell density (as per described in the “Cell density and live and dead assay” section) was also determined for each well.

Data were analysed on Graphpad Prism. Relative cell density (normalized by the density of Ctrl or lrECM untreated cells) was analysed by one way ANOVA, followed by multiple comparisons (all conditions *vs* Ctrl) using Two-stage linear step-up procedure of Benjamini, Krieger and Yekutieli (q<0.05 was considered significant). Differences in cell cycle based on the FUCCI probe expression were assessed by Two way ANOVA, followed by multiple comparisons (all conditions *vs* Ctrl, for G1, S/G2/M and just finished M/early G1) using Two-stage linear step-up procedure of Benjamini, Krieger and Yekutieli (q<0.05 was considered significant). These experiments were performed at least two times with two technical replicates (two wells/condition).

### Treatment with Pten inhibitor and analysis of downstream pathways activation

EpH4 cells were subjected to lrECM overlay assay with or without concomitant treatment with 2.5 µM bpV(pic) Pten inhibitor (control, Pteni, lrECM, lrECM+Pteni). 24h post treatment, cells were harvested for protein extraction and western blot analysis of protein expression. Data were analysed on GraphPad Prism using one way ANOVA followed by Tukey’s multiple comparisons test. These experiments were performed at least three times.

### Immunofluorescence of murine mammary tissue sections

The mammary fat pads were harvested from 5-weeks old female Balb/C mice obtained from Rede de Bioterios – USP, and the experimental procedures involved in this work were approved by the Ethics Committee for Animal Welfare from the Institute of Chemistry/USP (ethics number #218/2022).

Mammary fat pads were embedded in Tissue-Tek O.C.T., snap-frozen and stored at - 80°C. The samples were sliced at a thickness of 25 µm in a cryostat. Slices were collected in Starfrost slides, dried briefly and were stored at −80°C until used.

For immunofluorescence staining, slices were thawed, fixed in PFA 4% for 10 min and quenched in PBS-glycine 0.3 M. Antigen retrieval was performed by applying boiling Tris-EDTA (10 mM Tris base, 1 mM EDTA solution, pH 9) directly into the slides thrice with 5 min interval. Slices were then permeabilized in PBS-T 0.5% for 20 min, blocked in undiluted Odyssey Blocking buffer (LI-COR)/10% goat serum for 1h at room temperature and incubated with primary antibodies (anti-Pten, anti-Laminin-111, anti-IgG rabbit – table 1) overnight at 4°C. In the following day, slides were washed and incubated with fluorescence-conjugated secondary antibodies (table 2) and, when pertinent, with Phalloidin-Alexa 647 (1:50) for 45min at room temperature. Slides were counterstained with DAPI (0.5 µg/mL), mounted with Prolong Diamond antifade reagent, sealed and left O/N 4°C to cure.

### Mammary tissue sections imaging and analysis

Epithelial structures of interest (terminal end buds and ducts) were either imaged in Leica DMi8 wide-field fluorescence microscope (e-signal lab/USP) or in a Zeiss LSM880 Airyscan (INFABIC/UNICAMP). Several ducts and TEBs from mammary glands obtained from at least three mice were used for these analyses.

Images obtained in the Leica microscope were taken using the 63x/1.4NA oil objective in z-stacks and adjacent fields (to cover larger structures comprising whole ducts and/or TEBs) using the LasX Navigator Application (Leica microsystems) in three channels (DAPI for nuclei, FITC for Pten and TXR for Laminin-111). Images were initially processed in the LasX software, where adjacent fields were merged and 3D-blind deconvoluted. Five sequential Z-stacks were then combined to obtain the maximal projection. The resulting images were used to analyse the association between laminin content and Pten levels in ducts and TEBs using a customized pipeline on StrataQuest software (TissueGnostics). Details on parameters used for nuclei segmentation and detection on StrataQuest can be found at Russo *et al*., 2021^91^ and for this specific analysis, nuclei detection and segmentation was manually corrected for each image. In brief, images of ducts or TEBs were analysed separately; nuclei were detected in the DAPI channel and the mean intensity of Pten signal was obtained for each cell in an area comprised by the nucleus and the cytoplasm, which was estimated by growing outward from the nuclear area until neighbouring areas touched each other or until the fluorescence signal of Pten reached background levels (a similar approach to estimate cell boundaries was used for ADP ribose signal by Russo *et al*., 2021^91^). Total Laminin-111 fluorescent intensity (sum of intensity) was obtained for the entire field of interest (epithelial structure) and this value was divided by the total number of cells detected in that field (mean laminin-111 fluorescence). The average of the mean fluorescence intensity of Pten was plotted against the associated mean laminin-111 intensity for that field on GraphPad Prism and data were analysed by linear regression and Spearman’s correlation. Differences between Pten intensity in ducts *vs* TEBs were analysed by unpaired t-test.

Images obtained in the Zeiss LSM880 Airyscan were taken using the 63x/1.4NA oil objective and fluorescence was captured in four channels using the following laser lines for excitation: 405 nm (DAPI), 488nm (Pten, Alexa Fluor 488), 543 nm (Laminin-111, Alexa Fluor 546) and 633 nm (Phalloidin, Alexa Fluor 647). The detection spectra were adjusted manually for each channel to avoid crosstalk between the different channels. Both confocal and zoomed (3.4 zoom factor) super-resolution (Airyscan) images were obtained for ducts and TEBs at different Z points, corresponding to the basal domain (closer to the laminin-rich basement membrane) and apical domain (closer to the lumen).

### Analysis of quiescence and polarity in 3D acini upon Pten inhibition

#### Proliferation, roundness and size

EpH4-ES-FUCCI cells (5x10^3^ cells/cm^2^) were plated into 24-well plates on top of a bedding of Matrigel as previously described^20^. Wells were treated with 2.5 µM with the Pten inhibitor bpV(Pic) either on the day of seeding (day 0 and again at day 2 and day 4) or when quiescent 3D structures were already stablished (day 4). On day 5, 3D structures were incubated with Hoechst and imaged in Z stacks using a 10x objective in a fluorescent microscope (mCherry/mCitrine/Hoechst). Representative structures were then imaged at 40x. Image analysis was conducted using the LASX application (Leica). Briefly, images were deconvoluted and the maximum projection image was obtained. Structure size and roundness (0-1, being 1 a perfect circle) were obtained for individual structures, based on a mask built using the Hoechst channel. FUCCI-based cell cycle was calculated according to the mCherry and mCitrine areas in each field. These experiments were performed three times in duplicate wells and 3 fields/well at 10x were used for image analysis. Data were plotted in GraphPad Prism and analysed by one way ANOVA, followed by Tukey’s multiple comparisons test between all groups (Control, Pten inhibitor at day 0, and Pten inhibitor at day 4).

#### Onset and maintenance of Apico-basal polarity and lumen

EpH4 cells were seeded in low-adherence polyHEMA-coated plates (30000 cells/cm^2^), incubated for 48h to allow for cell clusters to be formed and replated in media supplemented or not with 4% of matrigel^26^. When indicated, cell clusters were treated with 2.5 µM with the Pten inhibitor bpV(Pic) either on the day of seeding (day 0) or when mature acini were already present (day 5). On day 6, 3D structures were fixed and subjected to immunofluorescence staining for ZO-1 (apical marker), α6 integrin (basal marker), Ki-67 (proliferation marker) and counterstained with DAPI (nuclei). Briefly, structures were fixed for 10 min by adding 32% PFA straight into the culture well (for a final concentration of 4%). Structures were then collected into microcentrifuge tubes, spun, resuspended and washed twice in 0.3 M PBS-glycine (quenching), permeabilized with 0.5% Triton-X in PBS for 20 min and blocked in 10% goat serum in Odyssey blocking buffer. Structures were then incubated overnight with primary antibodies diluted into Odyssey blocking buffer, washed thrice with PBS and incubated with Alexa Fluor conjugated secondary antibodies for 45 min (details on primary and secondary antibodies can be found in Tables 2 and 3), washed twice in PBS, incubated with DAPI for 5 min and mounted into glass slides using Prolong Diamond. All centrifugation steps were performed at 0.2 rcf for 3 min and all tubes and tips used were previously washed with 5% BSA in PBS. These experiments were performed in two biological replicates. Proliferation was estimated by the ratio between the area of Ki-67 positive cells and total area (DAPI). Slides were scanned in a Leica Dmi8 wide field microscope at 10x/0.3 NA (200 fields per slide), using the LASX Navigator application (Leica). Image analysis was conducted using the LASX application (Leica), where the total area for Ki-67 and DAPI were obtained and used to calculate the ratio between Ki-67 (proliferative cells) and DAPI (all cells) for a given slide.

The presence of lumen was assessed by manually scanning and counting the number of structures with lumen and the total number structures, using a 40x/0.6 NA objective in a Leica Dmi8 widefield microscope. The percentage of structures with lumen was then calculated for ach condition. At least 600 structures per slide were considered for these analyses.

The data regarding proliferation and lumen was then plotted and analysed in GraphPad Prism using ANOVA followed by Dunnett’s multiple comparisons test.

Representative structures were imaged on a Leica SP8-STED-FALCO microscope using the confocal mode, a 63x/1.4 NA oil objective and using the following laser lines for excitation: 405 nm (DAPI), 488 nm (Ki67, Alexa Fluor 488), 561 nm (integrin α6, Alexa Fluor 546), 633 nm (ZO-1, Alexa Fluor 647). The detection range for each emission spectrum was adjusted manually to prevent signal overlap.

## Acknowledgements

The authors would like to thank Hugo Armelin (Institute of Chemistry/USP, Brazil), Patricia Gama (Institute of Biological Sciences/USP, Brazil), William Festucchia (Institute of Biological Sciences/USP, Brazil), Cyrus Ghajar (Fred Huchinson Cancer Center, USA) and Sabrina Spencer (University of Colorado Boulder, USA) for the kind donation of antibodies, reagents and plasmids. We thank for the access to equipment and assistance provided by the National Institute of Science and Technology on Photonics Applied to Cell Biology (INFABIC) at the State University of Campinas; INFABIC is co-funded by FAPESP (#2014/50938-8) and CNPq (#465699/2014-6). The authors are also grateful to Nicolas Hoch and Deborah Schechtman for scientific discussions, advice on image analysis and insightful inputs.

